# Circulating tumor DNA profiling approach based on in silico background elimination guides patient classification of multiple cancers

**DOI:** 10.1101/2022.08.04.502459

**Authors:** Ming Li, Sisi Xie, Tao Hou, Tong Shao, Jingyu Kuang, Chenyu Lu, Xianling Liu, Lingyun Zhu, Lvyun Zhu

## Abstract

Circulating tumor DNA (ctDNA) analysis is increasingly providing a promising minimally invasive alternative to tissue biopsies in precision oncology. However, current methods of ctDNA mutation profiling have limited resolution because of the high background noise and false-positive rate caused by benign variants in plasma cell-free DNA (cfDNA), majorly generated during clonal hematopoiesis. Although personalized parallel white blood cell (WBC) genome sequencing suppresses the noise of clonal hematopoiesis variances observed in ctDNA based liquid biopsy, the system cost and complexity restrict its extensive application in clinical settings. To address this challenge, we describe a <u>Ma</u>tched WBC <u>G</u>enome sequencing <u>I</u>ndependent <u>C</u>tDNA profiling (MaGIC) approach, which enables the sensitive detection of recurrent tumor mutant information harbored by ctDNA from a bulk cfDNA background based on hybrid capture cfDNA deep sequencing, *in silico* background elimination, and a reliable readout measurement by the calculation the number of key mutated exons. Leveraging somatic mutation data from 10163 patients across 24 cancer types obtained from The Cancer Genome Atlas, we confirm that the MaGIC approaches are of ideal performance in prediction of prognosis by tissue biopsy samples of patients across multiple cancers. Meanwhile, MaGIC approaches enable the classification of prostate cancer patients from heathy cohorts by ctDNA sequencing data. We further profiled the ctDNAs of 80 plasma samples from 40 patients with nasopharyngeal carcinoma before and during chemotherapy by MaGIC approaches. The MaGICv2 can predict the chemosensitivity with high accuracy by simply using one liquid biopsy sample of each patient before a stereotypical treatment course. We anticipate that this new approach has the potential utility of ctDNA detection in multiple clinical cancer contexts, thus facilitating precise cancer therapy.

**Teaser:** A liquid biopsy analysis method for multifunctional patient classification such as diagnosis and chemosensitivity prediction.

## Introduction

Liquid biopsy, a minimally invasive and repeatable method for early diagnosis, prognosis prediction, and screening of cancers based on circulating biomarkers, has considerable clinical implications ^1^. The abundance of pre-clinical research has shown that circulating tumor DNA (ctDNA) released from apoptotic or necrotic tumor cells can act as an accessible biomarker detected by liquid biopsy ^2^. ctDNA can be detected in the blood of patients with multiple advanced cancers, and records valuable genetic information of the tumor genome, such as mutation signals and methylation profiles ^3^. Indeed, the blood tumor mutation burden (bTMB) and mean variant allele frequency (mVAF) measured by ctDNA sequencing represents the mutation density and abundance of the tumor genome, which can predict the outcomes of cancer immunotherapy and targeted therapy immunotherapy ^4,5^.

However, ctDNA analysis has not yet become a standard tool in the clinical oncologist’s arsenal because of its modest sensitivity, low coverage of patients, and/or high cost using current ctDNA selection panels and background elimination methods ^6^. Moreover, previous clinical trials have shown that ctDNA could not be detected in more than 50% of patients who ultimately recurred ^7,8^. Given that the major source of cell-free DNA (cfDNA) present in the blood is generated from hematopoietic cells, whereas less than 1% is tumor-derived, enrichment of trace amounts of blood ctDNA signals from the intense background noise associated with clonal hematopoiesis should be beneficial for improving ctDNA detection sensitivity ^9-11^. Recent approaches have designed different panels covering commonly mutated driver genes or exons from tumors to characterize small amounts of ctDNA in large populations of cfDNA ^12-15^. In parallel, paired white blood cell (WBC) sequencing has been conducted to eliminate the misclassification of WBC-derived variants, rendering the detection of ctDNA alterations more accurate ^16,17^. However, the increased system cost and complexity challenge the further clinical use of this approach. Theoretically, an *in-silico* background elimination strategy could be cost-effective. Indeed, Newman et al. developed a computational strategy to polish the background noise by modeling position-specific errors in a training cohort of healthy donors to allow error suppression in independent samples ^18,19^. Greater efforts should be made to further characterize the stereotypical background errors, improve the detection accuracy, and highlight the clinical implications of the ctDNA biomarkers.

In this study, we devised a <u>Ma</u>tched WBC <u>G</u>enome sequencing <u>I</u>ndependent <u>C</u>tDNA profiling (MaGIC) method, which synergically integrated a ctDNA capturing panel (termed Enricher), *in silico* background elimination approach (termed Filter), and optimal measurement strategy (termed KME) for ctDNA detection and monitoring (Figure 1). The ctDNA Enricher followed by TMB/bTMB readout measurement, initially exemplified by MaGICv1, was designed to target driver mutations and recurrently mutated exons within the cancer of interest, and can be used to hybrid capture next-generation sequencing of ctDNA, which is simply analyzed by the number of mutations. To further improve the ability of the selected biomarkers to classify patients with higher accuracy, we designed MaGICv2, a version that optimized the panel by filtering the potential benign mutant exons collected from the whole-exon-sequencing data of healthy donors from the public database. Moreover, the filtered mutation exons were readout by a new KME calculation measurement, which is more reliable than bTMB to increase the signal-to-noise ratio in the context of plasma ctDNA sequencing. By applying both MaGICv1 and v2 approaches in tissue and liquid biopsies, they can effectively predict patient survival in multiple cancer types and classify the patients from the healthy cohorts. More importantly, the MaGICv2 can accurately predict the chemotherapeutic response in locally advanced nasopharyngeal carcinoma (NPC) at the early stage before the stereotypical treatment course independent of paired WBC or tissue biopsy sequencing. This may be of great benefit for precision medicine as the doctors can reconsider the necessity of the two-round chemotherapy in a relatively reliable and affordable manner. This synergic detection system may substantially advance the sensitivity and application scope of ctDNA detection independent of other paired genome sequencing, thereby increasing the use of ctDNA-based liquid biopsy.

**Figure 1.**
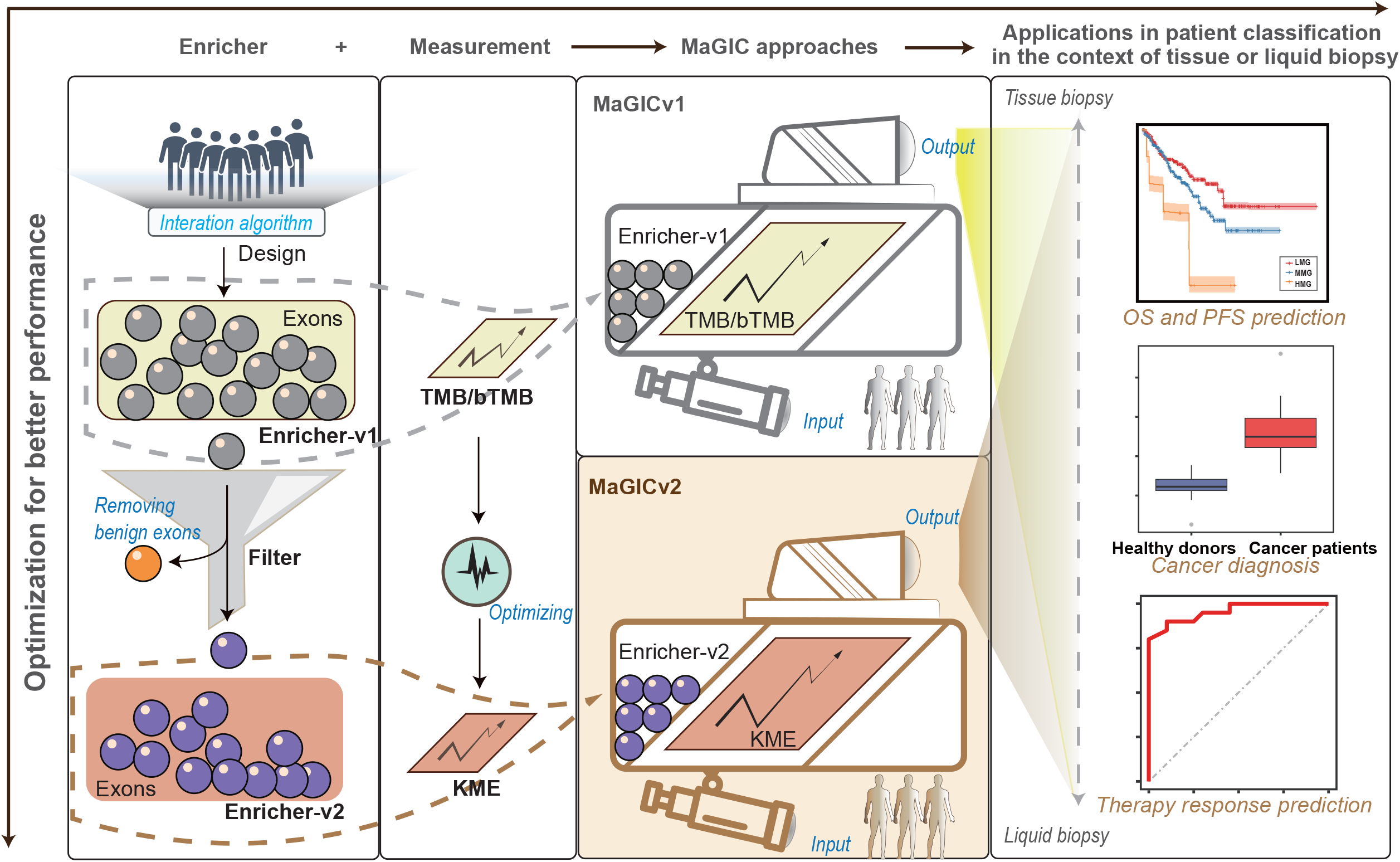
Schematics of the design and application of the MaGIC approaches. The MaGIC approaches were developed by systematical incorporation of the exon capturing panel with cancer-relevant mutations (Enrichers) and the effective mutation readout measurements (bTMB/TMB and KME). In the evolution of the two versions of MaGICs, a rational *in silico* background elimination process (Filter) was conducted to remove potential benign mutations that exist extensively in a healthy population. The MaGICs were sought to be applied in ctDNA detection for cancer diagnosis and, in particular, chemosensitivity prediction at a very early stage of the stereotype treatment course, as well as in prognosis evaluation in the context of tissue biopsies. ctDNA, circulating tumor DNA. TMB, tumor mutation burden. bTMB, blood-based TMB. KME, the number of key mutated exons. OS, overall survival. PFS, progress-free survival. LMG, low-mutation group. MMG, middle-mutation group. HMG, high-mutation group. ROC, receiver operating characteristic.

## Results

### Design of MaGICv1 for patient identification with high coverage

We leveraged an iterative algorithm to construct a minimal panel of genomic recurrent mutation exons that can capture most of the patients in different cancers with a repetitive detection frequency of each patient ^18^. To improve the capture rate of patients and initial input, the WES data processed by TCGA database, including 781 patients across TCGA-HNSC, TCGA-DLBC, and TCGA-SARC cohorts, were used as training datasets (Figure 2A). The measurement of the number of mutant exons (NME) was used to calculate the repetitive detection times, which reflect the detection robustness for each patient. The capture rate in each cohort was defined as the percentage of patients with NME > 0. After four sets of iterative calculations, 902 exons, with a total sequence length of approximately 610.7 kbp were selected to form Enricher-v1, which can capture 97.70% of patients at least once from the three pooled training datasets. Meanwhile, the capture rates of patients with more than two and three times of repetitive detection reached 85.28% and 71.57%, respectively (Figure 2B). The high capture rates were consistent in all three cancer datasets separately (Figure 2C). To test whether Enricher-v1 was available in multiple cancers, the WES data of 24 cancer types in TCGA were analyzed. As shown in Figure 2D, Enricher-v1 could identify > 90% of the patients across 11 cancer types, including esophageal, bladder, colorectal, lung, skin, prostate, stomach, ovarian, uterus, mesenchymal, and cervical, most of them comprising the top 10 most lethal cancers worldwide ^41^ (Supplementary Figure 1). These results indicate that Enricher-v1 is a potentially effective, robust, and versatile tool to identify various cancers by recurrent mutations. Considering the TMB as an emerging biomarker for patient stratification in oncology, we incorporated the TMB analysis into the Enricher-v1 mutation readout to form an analytical pipeline named MaGICv1.

**Figure 2.**
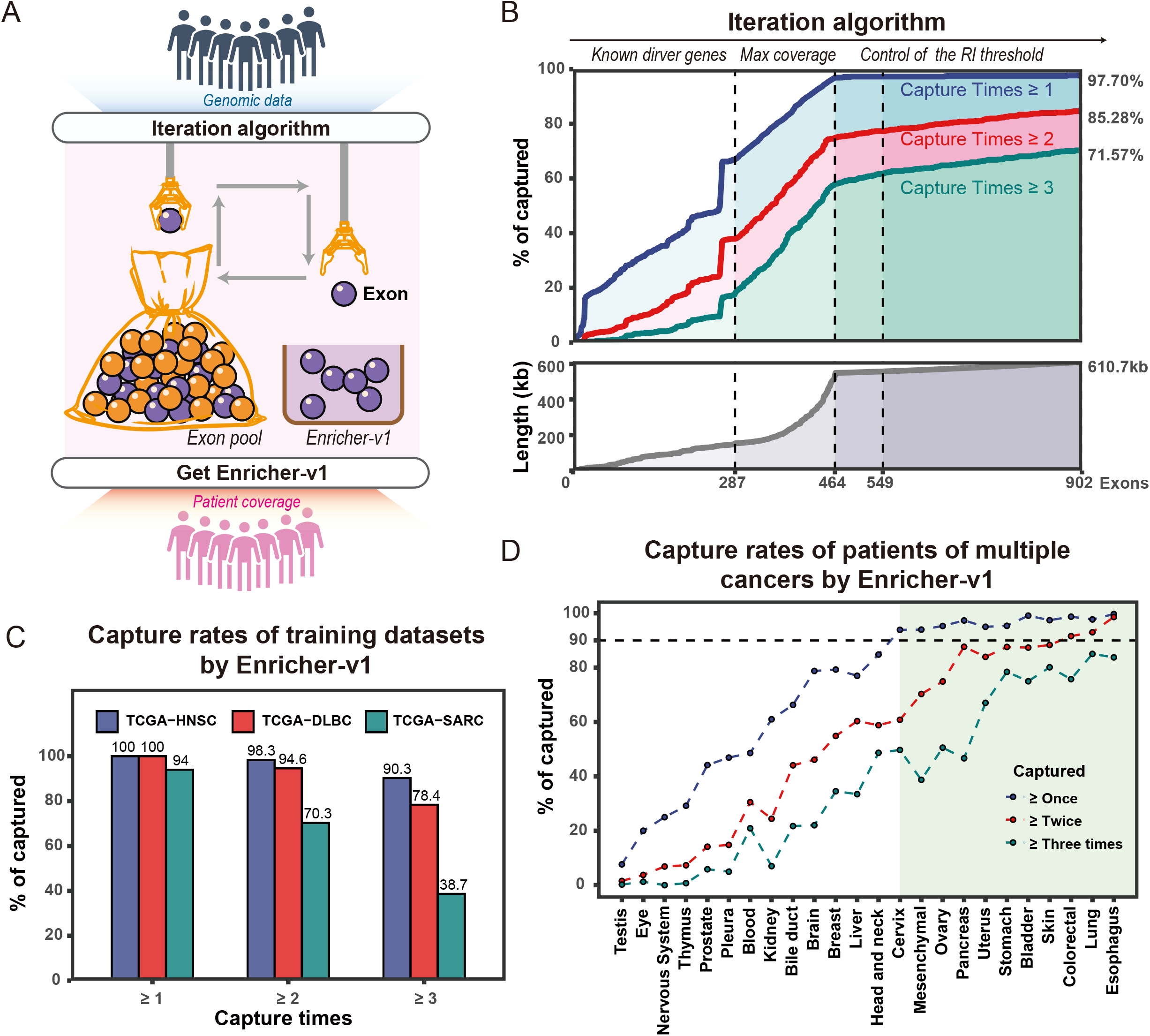
Design of MaGICv1. **a**. Schematics of the generation of Enricher-v1. The iteration algorithm was built to interrogate the genomic mutation data from the public cancer databases and rigorously select the exons enriched with cancer-relevant mutations from the human whole-exome pool. Enricher-v1 formed with the selected exons was used to identify the patients whose mutations occurred on the exon pool. **b**. The iteration algorithm phases and the patient coverage of Enricher-v1. The dashed lines indicate the number of selected exons at the end point of each iteration algorithm phase. The filled areas in blue, pink, and green indicate the proportion of patients in the training datasets that were captured at least once, twice, and three times, respectively, with Enricher-v1. **c**. Comparative analysis of the patient coverage of Enricher-v1 in three cancer datasets of training cohorts. **d**. The patient capture rate of Enricher-v1 in multiple cancer types. The cancer types with 90% of the patients who can be captured at least once with Enricher-v1 are highlighted in the green area. TCGA, the Cancer Genomic Atlas. RI, recurrent index. HNSC, head and neck squamous cell carcinoma. DLBC, lymphoid neoplasm diffuse large B-cell lymphoma. SARC, sarcoma.

### Performance of MaGICv1 in patient classification

To evaluate the ability of MaGICv1 in patient classification, we first tested its predictive value for OS and progress-free survival (PFS) based on the WES sequencing data of tumor tissue biopsies from 10163 patients with 24 different cancer types according to 33 TCGA projects (Supplementary Table 1). Through analyzing the TMB of the exons involved in Enricher-v1, the median and the upper quartile (Q3) values of MaGICv1 were 5 and 10, respectively, across 24 cancer types (Figure 3A). Considering the driver and recurrent mutations as being the major causes of tumorigenesis and progression ^42,43^, we envisioned that the patients with different mutation patterns would have different outcomes in prognosis, which may constitute an effective biomarker. Thus, we defined the patients with MaGICv1 TMB = 0 as the low-mutation group (LMG), the patients with MaGICv1 TMB > 5 (the median value of TMB) or > 10 (the Q3 value of TMB) as the high-mutation group (HMG), and the patients between the two as being the middle-mutation group (MMG). As expected, all three groups showed significantly different prognoses across various cancer types (Figure 3B and Supplementary Figure 2A). Notably, in several cases, such as kidney cancer and head and neck cancer, both significant PFS and OS benefits were observed within LMG groups, and significantly worse PFS and OS were found in HMG groups, suggesting that high mutation correlated with a high risk of poor prognosis (Figure 3C and Supplementary Figure 2B). Focusing on the top 10 most lethal cancers worldwide, more than 50% of them showed statistical discrepancies in the PFS and OS among all three HMG, MMG, and LMG groups (Supplementary Figure 3). These results suggest that MaGICv1-based WES sequencing with tissue biopsies is appropriate for the evaluation of prognosis in multiple cancers.

**Figure 3.**
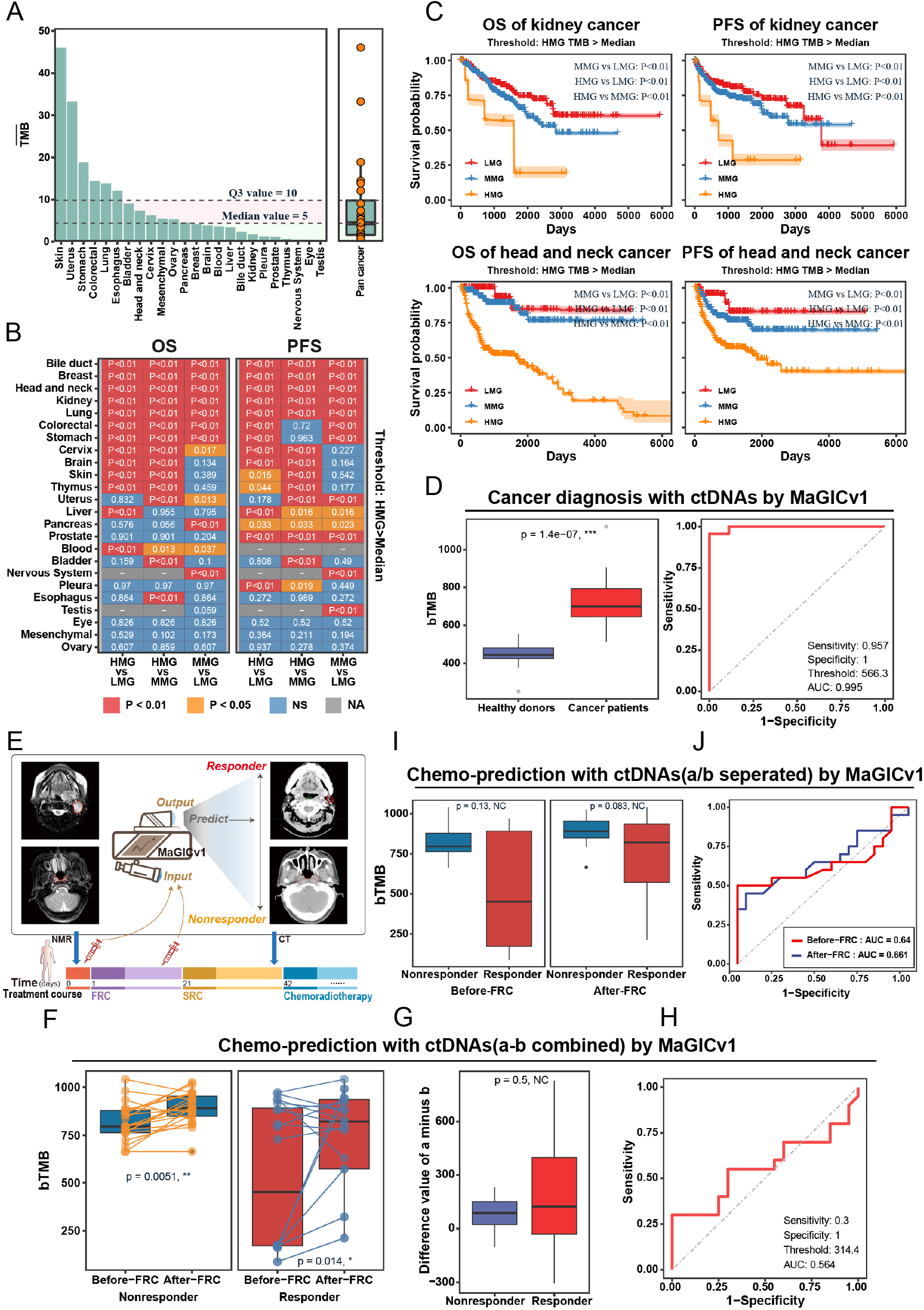
Performance of MaGICv1 in patient classification. **a**. Mean TMB 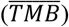 analysis of the exons in Enricher-v1 among various cancer types in TCGA. 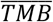 is defined as the average TMB of pateints in each cancer. The dashed line represents the Q3 and median values of the 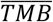 across all cancer types. The dot-boxplot indicates the distribution of 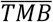 all datasets. **b**. Evaluation of the OS and PFS of patients with multiple cancers by MaGICv1. The patients are classified as HMG (TMB = 0), MMG (1 ≤ TMB ≤ median value threshold), and LMG (TMB > median value threshold) of each cancer. The P-value between each comparison is labeled in the heatmap and assigned to different colors (log-rank test). **c**. The univariable Cox curves of PFS and OS of representative cancer type (kidney cancer and head and neck cancer) where the patients are classified by MaGICv1 (HMG thresholds: median value). **d**. Classification of patients with cancer from healthy cohorts using ctDNA sequencing data by MaGICv1. **e**. Schematics of the application of MaGICv1 in chemosensitivity prediction. The photographs of responders and nonresponders are Patient R1 and Patient N9, as recorded in Supplementary Table 4. **f–h**. The performance of MaGICv1 in chemosensitivity prediction according to the ctDNA sequencing data of liquid biopsies before and after FRC. The difference level between responders and nonresponders to the chemotherapy is presented as paired bTMB analysis (**f**), bTMB difference analysis (**g**), and ROC analysis (**h**). **i, j**. The performance of MaGICv1 in chemosensitivity prediction according to the ctDNA sequencing data of separate liquid biopsy samples before or after FRC. The difference level between responders and nonresponders to the chemotherapy is presented as bTMB analysis (**i**) and ROC analysis (**j**). TMB, tumor mutation burden. bTMB, blood-based TMB. 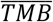, Mean TMB. Q3, the upper quantile. OS, overall survival. PFS, progress-free survival. HMG, high-mutation group. MMG, middle-mutation group. LMG, low-mutation group. TCGA, The Cancer Genome Atlas. NMR, nuclear magnetic resonance map. CT, computerized tomography. FRC, first-round chemotherapy. SRC, second round chemotherapy. ROC, receiver operation characteristic. AUC, area under curve. **P < 0.01, ***P < 0.001, NS, not significant. NA, data not available.

To further assess the patient classification outcomes of MaGICv1 in the context of liquid biopsies, we first used MaGICv1 to distinguish cohorts of healthy donors and patients with prostate cancer. The cfDNA WES data from patients with prostate cancer (BioProject ID: PRJNA554329) and healthy donors (SRA ID: SRP147273) were screened by Enricher-v1 and assessed by the bTMB. As shown in Figure 3D, the bTMB of the patients with cancer was significantly higher than that of the healthy donors (P = 1.4×10^−7^), and the ROC curves with an area under curves (AUCs) score of 0.995 revealed that the cancer population could be outstandingly distinguished from the healthy donors. Thus, this approach may provide an alternative means to perform minimally invasive diagnosis of prostate cancer.

Next, we collected 80 plasma samples from 40 patients with locally advanced NPC before- and after-the first-round chemotherapy (FRC) treatment and analyzed them by MaGICv1-based ctDNA capture sequencing to predict the therapeutic outcome via liquid biopsies (Figure 3E, Supplementary Figure 4 and 5). In both populations of the responders and nonresponders to chemotherapy, the bTMB of the Enricher-v1 panel in the after-FRC group significant increased compared with that in the before-FRC group (responders: p=0.014, nonresponders: p=0.0051). However, there was no significant difference between the responders and the nonresponders by calculating the bTMB difference data between the after- and the before-FRC (a-b) groups (P = 0.5; Figure 3G). The ROC analysis showed that the a-b is unable to group the responders and nonresponders with an AUC score of 0.564 (Figure 3H). Similarly, there was no significance between the bTMB of the responders and the nonresponders when analyzing the before- or after-FRC samples separately (P = 0.13 and 0.083, respectively; Figure 3I), and the AUC also showed poor scores (0.64 and 0.661, respectively; Figure 3J). These results suggest that MaGICv1-based ctDNA capture sequencing failed in prediction of chemosensitivity of patients with NPC using plasma samples at the early stage of a stereotypical treatment course. Further optimization of this approach is necessary to further reduce the background variants that may account for a large part of the ctDNA mutation signals readout by MaGICv1.

**Figure 4.**
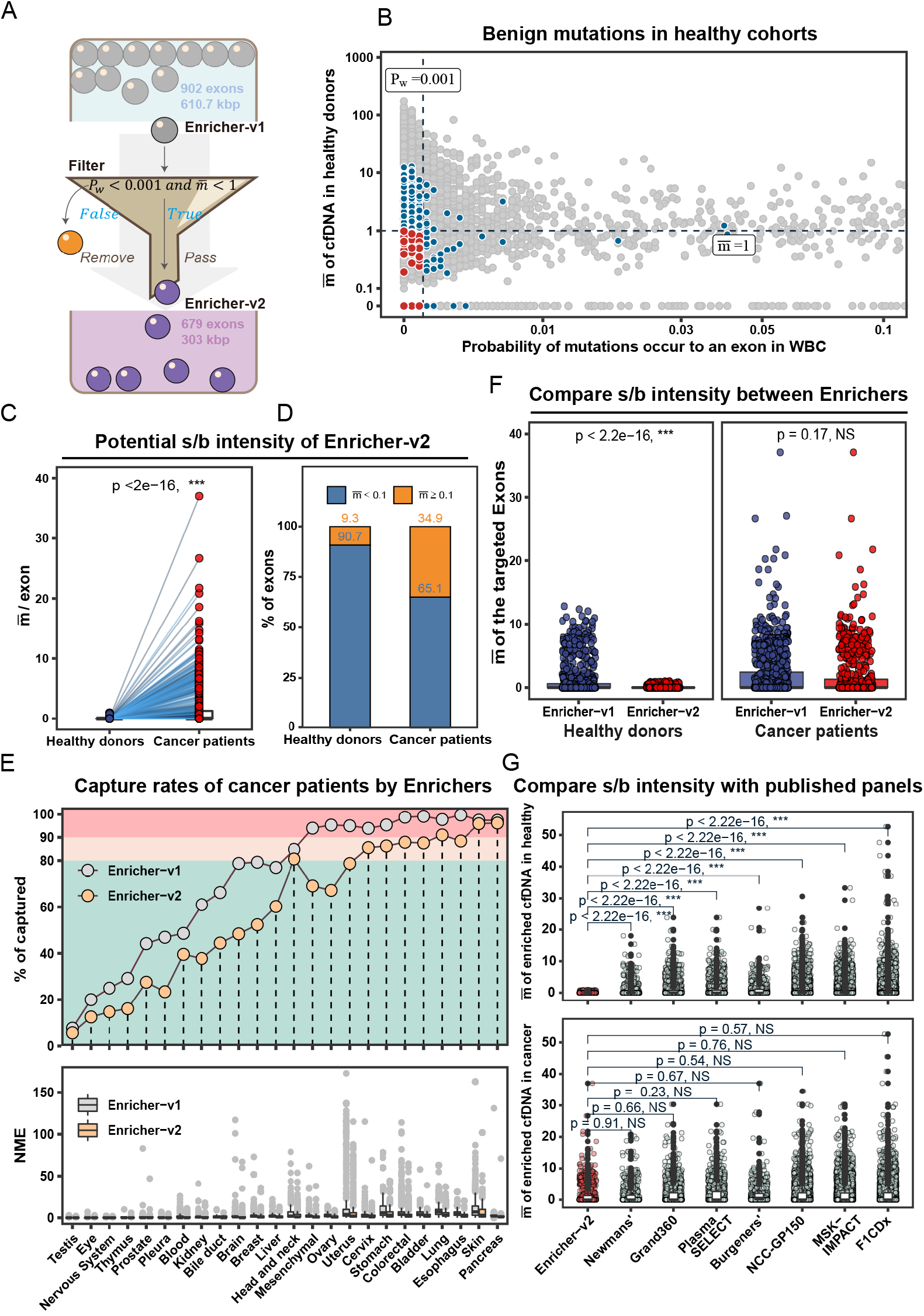
Development of Enricher-v2. **a**. Schematics of the workflow of Filter and the design of Enricher-v2. **b**. Acquirement of Enricher-v2 by filtering the exons with potential high benign mutation background in Enricher-v1. The gray dots indicate all of the exons in the entire human exome. The blue dots indicate the exons involved in Enricher-v1. The red dots indicate Enricher-v2 containing the filtered exons with low benign mutation frequency. **c, d**. The evaluation of potential signal to background (s/b) intensity of exons in Enricher-v2. The mutation features of patients with cancer and healthy cohorts are presented as paired analysis of the 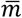 of each exon (**c**) and the percentage of exons with different mutation features (**d**) in Enricher-v2. **e**. Comparison of patient coverage between the two versions of Enrichers in multiple cancer types. The cancer types with 90% and 80% of the patients who can be captured by at least once with Enricher-v1 are highlighted in the pink and yellow areas. The boxplot shows the NME value of Enrichers across various cancer types. **f**. Comparison of s/b intensity between the two versions of Enrichers. **g**. Comparison of the s/b intensity of Enricher-v2 with that of other published genomic region panels for enriched cfDNA analysis. cfDNA, cell-free DNA. *P*_w_, mutant probability in healthy donors. 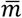, the average mutation frequencies per kilobase in healthy donors. s/b intensity, signal/background intensity. NME, the number of mutated exons. **P < 0.01, ***P < 0.001, NS, not significant.

**Figure 5.**
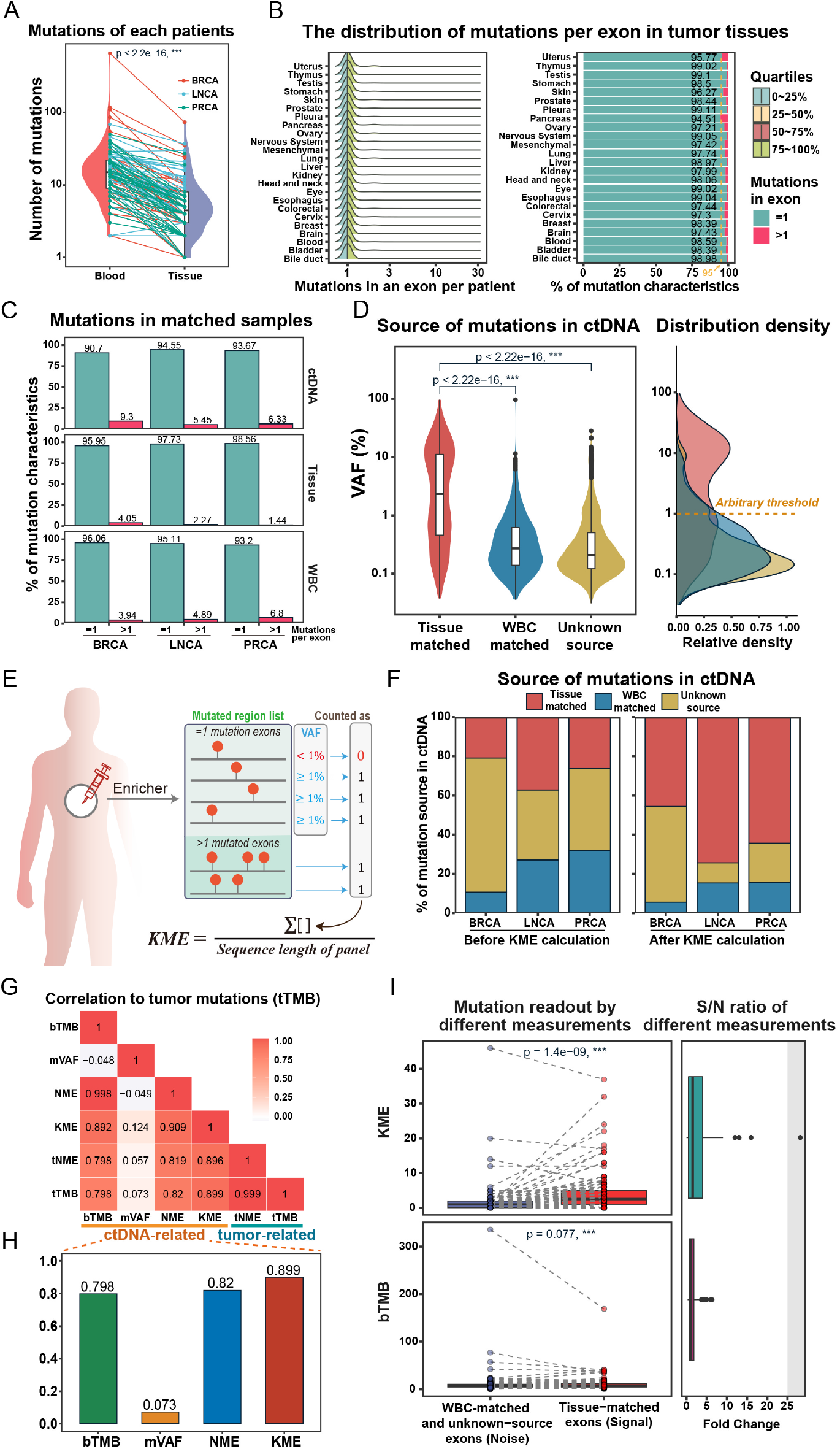
Design of KME measurement for MaGIC mutation readout. **a**. Analysis of the mutations of each patient harbored by the exons of Enricher-v2 in paired ctDNA and tumor tissue. The high-intensity sequencing data of matched ctDNA, WBC, and tumor tissues from 124 patients with metastatic cancer were used. **b**. The distribution of the exons of Enricher-v2 with different mutational features in tumor tissues across various cancer types. All of the mutations in the TCGA datasets were included. **c**. The proportion of exons of Enricher-v2 with different mutational features in matched sequencing data of ctDNA, WBC, and tumor tissues from patients with breast cancer (BRCA), lung cancer (LNCA), and prostate cancer (PRCA). **d**. The variant allele frequencies (VAFs) of the ctDNA mutations in matched WBC, tumor tissues, or unknown sources. A VAF = 1% was arbitrarily set as the threshold to define the ctDNA mutation with or without high confidence to reflect the mutation in tumor tissue. **e**. Schematics of the calculation of KME measurement. **f**. Comparison of the proportion of effective ctDNA mutations in Enricher-v2 matched to different sources before and after KME calculation. **g, h**. Pearson correlation analysis of different ctDNA-related and tumor-related measurements. The bar chart shows the Pearson correlation coefficients of ctDNA-related measurements with tTMB. **i**. Comparison of the signal-to-noise (S/N) ratio of ctDNA-related measurements according to the paired values (the left boxplot) and fold changes (the right boxplot). BRCA, breast cancer. LNCA, lung cancer. PRCA, prostate cancer. VAF, variant allele frequency. KME, the number of key mutation exons. mVAF, mean VAF. bTMB, blood-based tumor mutation burden. tTMB, tissue-based tumor mutation burden. NME, the number of mutation exons. tNME, the NME calculated by tissue samples. cfDNA, cell-free DNA. ctDNA, circulating tumor DNA. S/N ratio, signal-to-noise ratio. ***P < 0.001, NS, not significant.

### Background suppression of Enricher-v1 for MaGIC optimization

Inspired by the recognition that biological confounding factors generated by the hematopoietic cells seriously challenge the sensitivity of ctDNA detection ^44^, we sought to improve the performance of MaGIC-based ctDNA analysis for clinical patient classification by suppressing these potential benign mutant backgrounds in the Enricher-v1 panel. As previously described, hematopoietic cells accumulate clonal hematopoiesis variances (CHV) of indeterminate potential during aging and constantly release their genomic DNA segments into the blood to form a major component of the ctDNA pool ^45,46^. A matched cfDNA and WBC sequencing approach has been shown to provide a promising resolution in differentiating the mutation signal of tumors from the variants related to clonal hematopoiesis ^17^. To develop an alternative efficient strategy that is both simpler and cheaper for the patients, we designed an *in silico*-based Filter by data mining the noisy mutations from both WBC and cfDNA sequencing information of a large population of healthy individuals. The process of the Filter started with the ctDNA Enricher-v1 exon panel and was performed in two stringent steps: (i) remove the exons with the mutant probability (*Pw*) > 0.001 in the WBC of healthy individuals; and (ii) remove the regions with the average benign mutation frequencies per kilobase 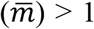 in the cfDNA of healthy individuals (Figure 4A, Methods). For step i, the WES sequencing data of the WBC of 2504 healthy individuals from the 1000 Genome Project database were analyzed ^36^. For step ii, the cfDNA sequencing data from nine healthy donors were analyzed ^21^. After filtering, 679 exons covering 303 kbp sequences (hereinafter referred to as Enricher-v2) were selected from the beginning of the 902 exons covering the 610.7 kbp sequence in the Enricher-v1 panel (Figure 4B).

To evaluate the potential signal-to-background (s/b) intensity of Enricher-v2 in the ctDNA analysis, the cfDNA sequencing data from patients with prostate cancer (BioProject ID: PRJNA554329) ^22^ and healthy donors (SRA ID: SRP147273) ^21^ were analyzed. The 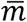 of each exon involved in Enricher-v2 was calculated in these datasets. The exons in Enricher-v2 had significantly higher 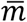 in the cfDNA of the patients (signal 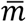) than of the healthy individuals (background 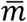) (Figure 4C). There were also displayed striking differences in the distribution of 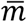 in these two groups, where 90.7% of exons in the healthy donors had a background 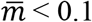, while 34.9% of exons in the patients with cancer had a signal 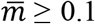 (Figure 4D). Compared to Enricher-v1, the rate of capturing patients with cancer at least once were not obviously decreased using the size-reduced Enricher-v2 panel (Figure 4E), whereas a better s/b intensity was acquired since Enricher-v2 showed a comparable signal 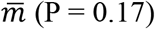 but significantly lower background 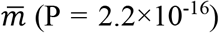 (Figure 4F). Upon further comparison with the recently published Enricher panels for ctDNA detection ^18,23-28^, Enricher-v2 showed significantly higher potential s/b intensity and lower false-negative detection rates, with comparable patient capture rates (Figure 4G). Thus, the Filter not only eliminated potential benign mutants but also reduced the size of the Enricher, which may further improve the detection sensitivity and simplify the analytical process of the MaGIC approach.

### Design of KME measurement for MaGIC optimization

We next envisioned that using bTMB for the indiscriminate measurement of the tumor-derived somatic mutations in the ctDNA pool may record significant background variants and result in false-positive identification. To confirm this hypothesis, we analyzed the high-intensity sequencing data of captured ctDNA, matched WBC, and tumor tissue biopsies from 124 patients with metastatic cancer presented in a previous rigorous study ^20^. The detection frequency of mutations in the ctDNA was significantly higher than that in the matched tumor biopsies, indicating that a large proportion of the mutation signals were not derived from the tumors (Figure 5A). Interestingly, more than 95% of the mutant exons in the genomes of different tumors only carried one mutation according to TCGA mutation datasets (Figure 5B), and similar results were observed in the analysis of matched sequencing of the ctDNA and the tumor tissue biopsies, indicating that the exons with more-than-one mutations might contain background variants not from tumor source (Figure 5C). To interrogate the origin of the ctDNA mutation repertoire, we mapped the mutations in the one-mutation exons of the ctDNA to the matched sequencing data of the WBC and tumor biopsies and calculated the variant allele frequency (VAF) that is the percentage of sequence reads observed matching a mutation divided by the overall coverage at that locus. The distribution of the VAF differed significantly between the tumor-matched mutations and the potential background mutations (WBC-matched and unknown source), and a VAF of 1% could be set as a threshold to remove most of the backgrounds while retaining sufficient tumor mutation signals (Figure 5D). Encouraged by these observations, we sought to define a new measure, termed the number of key mutation exons (KME), to improve the accuracy of the identification of tumor-derived somatic variants from ctDNA (Figure 5E).

From this measurement, the ctDNA mutation data acquired by the Enricher-based capture sequencing were counted by the NME to avoid interference of the more-than-one mutation exons. Subsequently, the mutations with VAF < 1% were removed, and the remaining mutation-harboring exons were analyzed. Indeed, with the measurement of KME, the mutations of ctDNA revealed the tumor-derived signals more reliably (Figure 5F). The Pearson correlation coefficient analysis also showed a stronger correlation of the tumor-derived TMB (tTMB) with the KME (r = 0.899) than with other conventional measurements, such as the mVAF (r = 0.073), bTMB (r = 0.798), and NME (r = 0.820) (Figure 5G, H). The signal-to-noise ratio, calculated as the number of tumor tissue-matched exons divided by that of the WBC-matched and unknown-source exons, was remarkably increased with the KME measurement (Figure 5I). Therefore, the KME may be a valuable measurement in the MaGIC approach to mirror the ctDNA mutations from the tumor-derived signals with an improved signal-to-noise ratio.

### Performance of MaGICv2 in the context of tissue biopsies

By incorporating the optimized Enricher-v2 and the KME measurement into MaGIC, MaGICv2 was devised. We then tested the ability of MaGICv2 to predict the OS and PFS of patients with multiple TCGA cancer types (Figure 6A). Similar to the above circumscription, the LMG, HMG, and MMG were defined to include the patients with MaGICv2 KME = 0, MaGICv2 KME > threshold (median value of 5 or Q3 value of 10), and the MaGICv2 KME between them, respectively. The MaGICv2 measured using TMB and mVAF was also analyzed by the same algorithm for comparison (Figure 6B). By statistical analysis, KME showed comparable or, in some cases, better performance than the TMB and mVAF in the classification of patients with a different OS or PFS, irrespective of the median value or at the Q3 value in the setting of the threshold (Figure 6C, Supplementary Figure 6A and Supplementary Figure 7). In particular, the KME showed remarkable significance in the classification of patients with breast cancer and kidney cancer (Figure 6D and Supplementary Figure 6B). As expected, both MaGICv2 and MaGICv1 have good performance in the prediction of the OS and PFS in multiple cancer types (Figure 6E and Supplementary Figure 8), while MaGICv2 outperformed MaGICv1 in prognosis prediction of the world top 10 most lethal cancer types (Supplementary Figure 9). Moreover, in comparison with other published ctDNA Enricher panels along with different measurements ^18,23-28^, MaGICv2 enabled efficient classification of patients with different OS and PFS in more cancer types, especially in the top 10 most lethal cancers worldwide (Supplementary Figure 10). Collectively, these data suggest that MaGICv2 may be an approach with high accuracy to predict the clinical outcomes of patients with different cancer types via tumor tissue biopsies.

**Figure 6.**
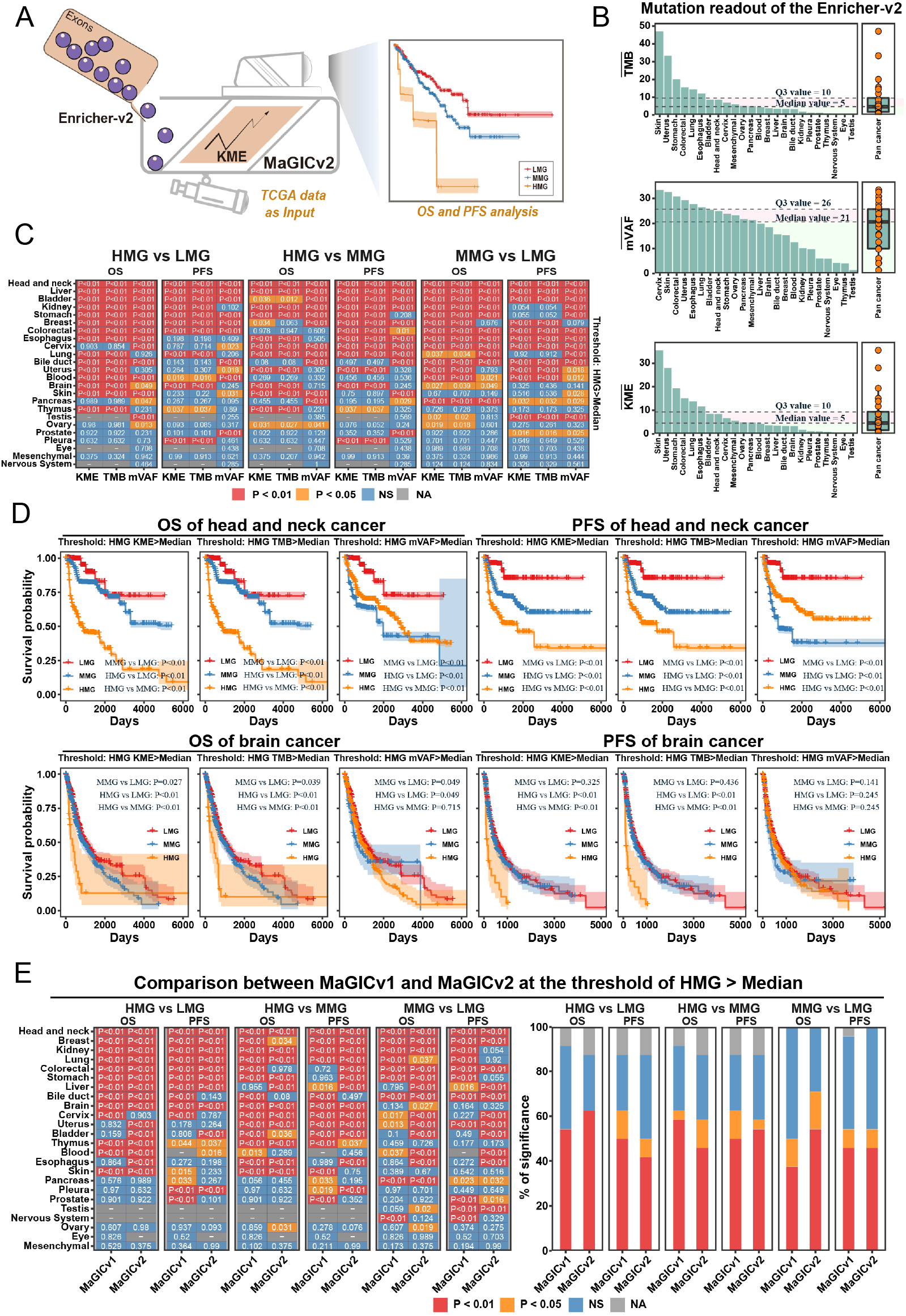
Performance of MaGICv2 in prognosis evaluation in the context of tissue biopsy. **a**. Schematics of the workflow of MaGICv2 for prognosis evaluation. **b**. Different measurements (TMB, mVAF and KME) for mutation readout in Enricher-v2 among various cancer types in TCGA. The dashed line represents the Q3 and median values of the readout data across all cancer types. The dot-boxplot indicates the distribution of 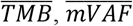 and 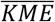 for all datasets. **c**. Evaluation of the OS and PFS of patients with multiple cancers by MaGICv2. The patients are classified as HMG (KME = 0), MMG (1 ≤ KME ≤ median value threshold), and LMG (KME > median value threshold) of each cancer. The P-value between each comparison is labeled in the heatmap and assigned to different colors (log-rank test). **d**. The univariable Cox curves of PFS and OS of representative cancer types (head and neck cancer and brain cancer) where the patients are classified by MaGICv2 (HMG thresholds: median value). **e**. A comparison of the performance of the two versions of MaGICs in prognosis evaluation (HMG thresholds: median value). Q3, the upper quantile value. OS, overall survival. PFS, progress-free survival. LMG, low-mutation group. MMG, middle-mutation group. HMG, high-mutation group. TMB, tumor mutation burden. KME, the number of key mutant exon. mVAF, mean variance allele frequency. 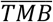, mean TMB. 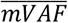, mean VAF. 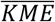, mean KME. NS, not significant. NA, data not available.

**Figure 7.**
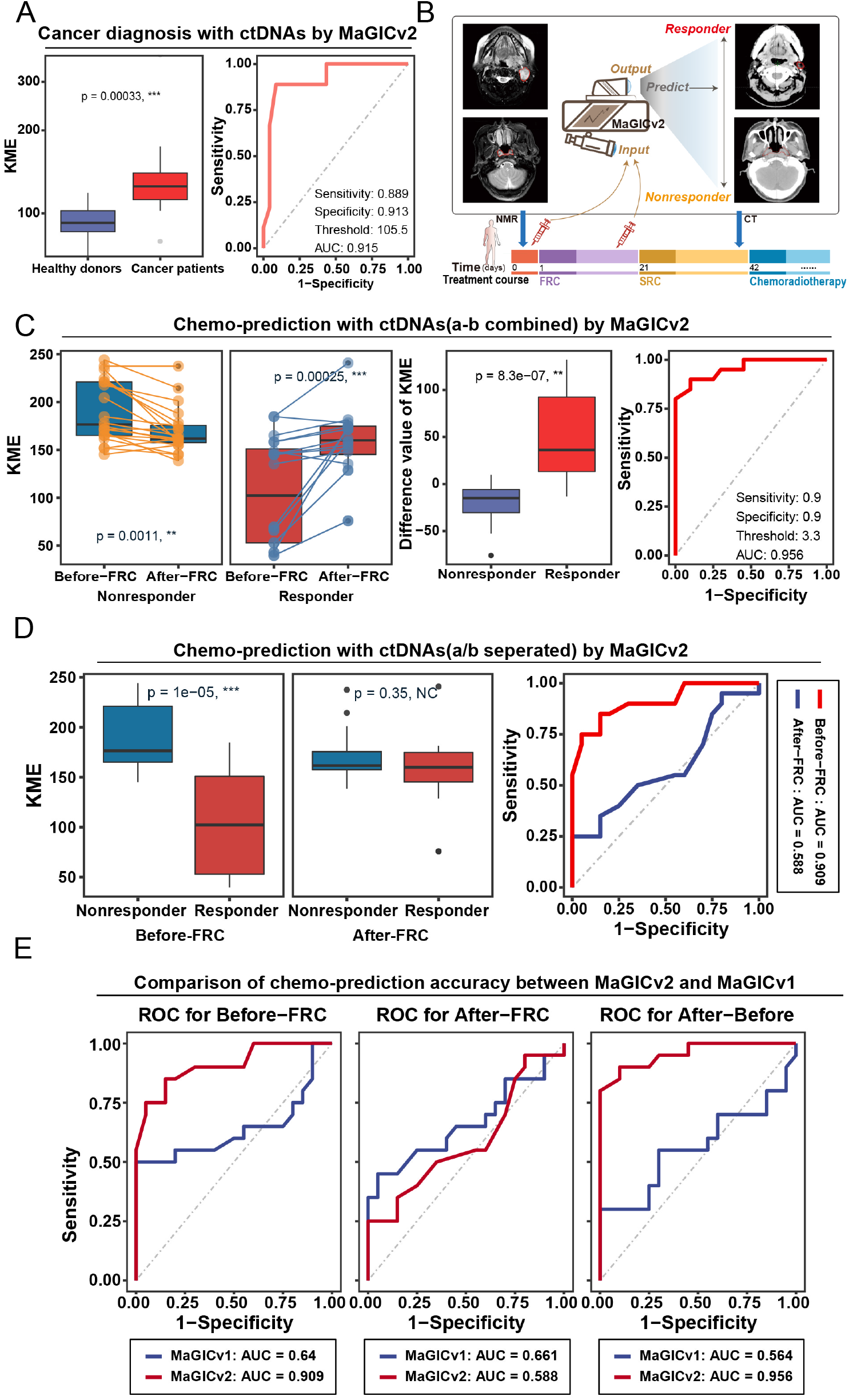
Performance of MaGICv2 in patient diagnosis and chemosensitivity prediction in the context of liquid biopsy. **a**. Classification of patients with cancer from healthy cohorts using ctDNA sequencing data by MaGICv2. **b**. Schematics of the application of MaGICv2 in chemosensitivity prediction. The photographs of responders and nonresponders are Patient R1 and Patient N9, as recorded in Supplementary Table 4. **c**. The performance of MaGICv2 in chemosensitivity prediction according to the ctDNA sequencing data of both the liquid biopsies before and after FRC. The difference level between responders and nonresponders to the chemotherapy is presented as paired bTMB analysis (the left graph), bTMB difference analysis (the middle graph), and ROC analysis (the right graph). **d**. The performance of MaGICv2 in chemosensitivity prediction according to the ctDNA sequencing data of separate liquid biopsy samples before or after FRC. The difference level between responders and nonresponders to the chemotherapy is presented as bTMB analysis (the left graph) and ROC analysis (the right graph). **e**. Comparison of the performance of the two versions of MaGICs in chemosensitivity prediction using single or both liquid biopsy samples. NPC, nasopharyngeal carcinoma. KME, the number of key mutation exons. ROC, receiver operation characteristic. AUC, the area under curve. FRC, first-round chemotherapy. mVAF, mean value of variant allele frequency. KME, number of key mutated exons. TMB, tumor mutation burden. bTMB, blood-based TMB. *P < 0.05, **P < 0.01, ***P < 0.001, NS, not significant.

### Performance of MaGICv2 in the context of liquid biopsies

We further tested MaGICv2 in the analysis of ctDNA liquid biopsies, which may help to assist cancer diagnosis or monitor therapy sensitivity. Similar to MaGICv1, the cfDNA WES data from 23 patients with prostate cancer (BioProject ID: PRJNA554329) and 9 healthy donors (SRA ID: SRP147273) were analyzed by MaGICv2. As well as the MaGICv1, MaGICv2 showed good performance in classification of patients from the healthy cohorts. The cancer patients showed significantly higher KME than the healthy cohorts (p=0.00033), indicating higher tumor mutation signals recorded in plasma ctDNA. The ROC curve showed good AUCs score of 0.995 (Figure 7A). To test the application of MaGICv2 in another scenario of therapeutic effect prediction, we analyzed 80 plasma samples from 40 patients with locally advanced NPC before- and after-FRC treatment using MaGICv2-based ctDNA capture sequencing (Figure 7B). Through MaGICv2, the responders showed a significantly increased trend in the after-FRC samples compared to the before-FRC samples (p=0.00025), while the nonresponders showed decreased trend (p=0.0011, Figure 7C). These results may be reasoned by more potent tumor cell senescence and tumor DNA release during chemotherapy in the responders than that in the nonresponders. As expected, the results of a-b could outstandingly distinguish the responders and the nonresponders with p value of 8.3×10^−7^ and ROC AUC score of 0.956 (Figure 7C). Moreover, there was a significant differentiation between the responders and nonresponders according to their before-FRC samples (p =1.0×10^−5^, ROC AUC=0.909), indicating the advantage of MaGICv2 in predicting chemosensitivity via a single sample from each patient (Figure 7D). Comparing MaGICv2 with different combinations of Enrichers and measures (including the MaGICv1), we found that MaGICv2 consistently produced the most accurate discrimination between responders and nonresponders early during chemotherapy in NPC (Figure 7E and Supplementary Figure 11). These data collectively demonstrate that the ctDNA analysis based on MaGICv2 could effectively classify the patients with prostate cancer from healthy people and, more importantly, predict the chemosensitivity of patients with NPC as early as before the treatment course in a simplified and cost-effective manner with the sample collection and analysis performed once only.

## Discussion

Many studies of ctDNA-based liquid biopsies have designed sequencing panels that cover hotspot mutations of exons from key cancer genes. However, the ctDNA sequencing assays using these panel-probed hybridizations remain inaccurate to some extent because the mutation signal output of these assays includes many variants that are absent in the respective tumor tissues and are inferred to be somatic. A prospective strategy with joint high-intensity sequencing of both cfDNA and WBC could mitigate the mutation detection artifacts, but it inevitably resulted in a cost increase for patients upon clinical use. Here we report MaGIC, an approach that enables robust de novo detection of tumor-derived mutations out of the massive repertoire of somatic variant backgrounds based on single ctDNA capture sequencing and *in silico* background elimination. This approach offers many possible choices of patient classification, including diagnosis, prognosis, and prediction of therapeutic benefits with liquid or tissue biopsies across multiple cancer types.

MaGICv1 was initially designed with a new panel that could capture most of the patients across more than 10 cancer types. MaGICv1 can evaluate the prognosis of many cancer types via tissue biopsies and can classify the patients from the healthy cohorts of patients via liquid biopsies by interrogating the TMB/bTMB. However, it failed in prediction of chemosensitivity of patients with NPC at the early stage of a stereotypical treatment course. In MaGICv2, we optimized the panel by eliminating the exons with potential non-tumor-matched nonsynonymous somatic mutations according to the benign variants presented by both the tissue genomics and the cfDNA sequencing data of healthy people. The optimized panel, with a small size of only 679 exons covering 303 kbp sequences, is comparable to several other published panels but showed the highest signal-to-background intensity among them. We then devised the KME, which further removed the mutation signals with similar incidences to the potential backgrounds and were more likely to recapitulate the distribution feature of the mutations stemmed from tumors. The measurement of KME outperformed other conventional methods such as bTMB and mVAF, particularly in the context of ctDNA analysis, thus may supply a substitutive strategy for evaluating the genomic mutation density. The MaGICv2-based ctDNA capture sequencing and the following bioinformatics analysis shows comparable efficacy in prognosis prediction and diagnosis via tissue and liquid biopsies, respectively. Intriguingly, the MaGICv2 achieves high accuracy in prediction of chemosensitivity of patients with NPC independent of any paralleled sequencing. Given that the treatment course of locally advanced NPC often includes two rounds of chemotherapies, MaGICv2 may account for a simpler and more cost-effective way to predict the outcome after this schematized course by acquiring only one liquid biopsy sample of each patient as early as before the first-round chemotherapy. This method may help to save monitoring time and avoid overtreatment to advance precision medicine.

However, as the patient population in this study is limited in size and is not necessarily representative of the global target population, it is necessary to perform further testing on additional large patient cohorts within the context of clinical trials, which will provide a more accurate estimate of the performance of this approach. Moreover, much additional efforts using state-of-the-art technologies such as machine learning are needed to further improve the sensitivity of the ctDNA detection approach to predict the therapeutic outcome. Interfacing MaGIC with more cases of patient classification across different cancer types is essential to explore its potential to enable extensive clinical applications.

In summary, this study showcases the power of the MaGIC approach for ctDNA sequencing independently of any parallel WBC or tissue sequencing, thus enabling a high-accurate and relatively low-cost method for the clinical use of promising ctDNA biomarkers. Given its prevalent effectiveness in the context of both tissue and liquid biopsies in multiple cancer types, the MaGIC approach may have broad implications for patient stratification and clinical decision-making.

## Materials and Methods

### Acquirement of the public data

The public data used in this study were acquired via the public databases and previous literature. The public databases include The Cancer Genome Atlas (TCGA, https://cancergenome.nih.gov/), the 1000 Genomes Project (1000G, https://www.internationalgenome.org), and the Sequence Read Archive (SRA, https://www.ncbi.nlm.nih.gov/sra). The data of matched mutation calling lists among the cfDNA, WBC, and tissue biopsy from 124 patients with lung, prostate, and breast cancers were obtained from Razavi et al. ^20^. The whole-exome sequencing (WES) data of cfDNA in 9 healthy donors were generated by Teo et al. ^21^. The mutation calling list of ctDNA in 23 patients with prostate cancer was obtained from Ramesh et al ^22^. Several published ctDNA capturing panels were used to perform a comparison with the Enrichers designed in this study, including the CAPP-Seq-NSCLC ^18^, CAPP-Seq-HNSC ^23^, Grand360 ^24^, Plasma SELECT ^25^, NCC-GP150 ^26^, MSK-IMPACT ^27^, and F1CDx ^28^. The human whole-exome was acquired by the incorporation of the exome coordinates from GTF files (gencode.v22.annotation.gtf.gz). The mutation coordinates of the human whole-exome in all public datasets were converted to hg38 for consistency by the function import.chain of the R package rtracklayer. Genomic regions of the panels were extracted and mapped to the human whole-exome for the following comparative analyses.

### Design of Enricher-v1

Enricher-v1 was designed by the iteration of recurrence mutation exons among TCGA database, and the recurrent index (RI), defined as the number of unique patients with somatic mutations per kilobase of a given genomic unit ^18^, was used to assess the frequency of mutations occurring in an exon. Originally, the Mutation Annotation Format (MAF) files recording the genomic SNVs of head and neck squamous cell carcinoma, lymphoid neoplasm diffuse large B-cell lymphoma, and sarcoma, presented in TCGA-HNSC, TCGA-DLBC, and TCGA-SARC, respectively, were set as the primary datasets. The mutation exons in the MAF files were mapped to the human whole-exome to build the primary exon pool. Subsequently, we calculated the RI of every exon in the primary exon pool. All of the exons were ranked by the RI. The Enricher-v1 design algorithm was developed according to a previous publication with modifications ^18^. Briefly, the driver genes listed by the PANCANCER project in ICGC were used for the Phase 1 exon panel selection. In Phase 2, the mutation exons covering more than five patients in HNSC, DLBC, and SARC were listed as candidates. Each exon that helps to identify more than one additional patient who has never been covered can be further selected. The calculation was iterated in the exon candidates in descending order of their RI. Afterward, the remaining exons with an RI of more than 30 (Phase 3) or 20 (Phase 4) and with more than three patients’ coverage were picked up and were further used as candidates for the above iterative calculation to improve patient coverage. After the robustness and capture rate reached a certain threshold, the exon list, termed Enricher-v1, was formed (Supplementary Table 2).

### Design of the Filter and generation of Enricher-v2

To evaluate the exons that commonly carried benign mutations in the Enriche-v1, we set up two variables, including the mutant probability (*P*_*w*_) and the average mutation frequencies per kilobase 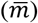 in healthy donors. The *P*_*w*_ was calculated to evaluate the probability of occurrence of potential clonal hematopoiesis variances (CHV) in WBCs according to the massive WBC mutation data from 2504 healthy donors. The mutation data were summarized and the frequency of each mutation occurring in the total donors was calculated. The mutations with frequencies > 1% were discarded to avoid the interference of potential single nucleotide polymorphisms (SNP). Subsequently, the remaining mutations were mapped to the human whole-exome, and the number of donors carrying each mutant exon was calculated. The formula of the *P*_*w*_ is as follows:

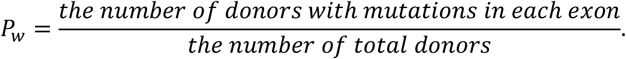

Meanwhile, the 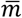 was calculated to remove the bulk background noises of the cfDNA using the published ctDNA sequencing data from healthy donors (SRA ID: SRP147273). First, the mutations in the dataset that passed the quality control (QC) were mapped to the whole human exome. The 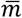 of each exon 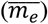 was defined as the sum of the number of mutations in this particular exon of all donors 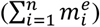 divided by the total number of donors (*n*) and the length of this exon (*l*_e_, kbp). The formula of the 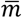 is as follows:

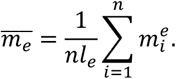

The exons with *P*_w_ ≥ 0.001 or 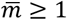 were filtered from Enricher-v1 to form the exon list of Enricher-v2 (Supplementary Table 3).

### Design of the measurements

We explored the performance of conventional clinically relevant measurements for mutation analysis, including the tumor mutation burden (TMB)/blood-based TMB (bTMB) and the mean variant allele frequency (mVAF), and compared them with our newly developed measurement, termed the number of key mutant exons (KME). The TMB/bTMB was defined as the number of somatic mutations (*m*) per kbp of the interrogated exons (*L*) with the following formula:

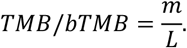

The mVAF was defined as the average number of variant reads divided by the number of total reads of the interrogated exons, with the following formula:

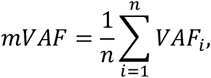

where the *VAF*_i_ represents the variant allele frequency (VAF) of mutation *i* and *n* represents the total number of mutations. To remove the potential CHV inference in the context of ctDNA detection, the KME calculation was started by filtering the exons that carried one mutation whose VAF was < 1%. The number of remaining mutant exons was directly counted (*R*) and then divided by the sequence length of these exons in kbp.

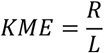

In the context of tissue biopsy analysis, the VAF filtering process was skipped owing to the inexistence of CHV inference in tumor tissues. The KME for the tissue biopsy could be defined directly as the total mutant exons per kbp sequence length of the exon list, which is also termed the number of mutant exons (NME).

### Collection of the plasma samples from patients with nasopharyngeal carcinoma

Plasma samples of patients with nasopharyngeal carcinoma were collected from the Department of Oncology, the Second Xiangya Hospital of Central South University, China. Informed consent was obtained after the nature and possible consequences of the studies were explained. All of the patients were treated with two cycles of chemotherapy followed by a concurrent chemoradiotherapy within three months after diagnosis. The nuclear magnetic resonance (NMR) imaging and the computerized tomography (CT) scans were examined at the time of primary diagnosis and after the second-round chemotherapy, respectively. The plasma samples of the patients upon the diagnosis and right after the first round of chemotherapy were used in this study. The patient characteristics are listed in Supplementary Table 4.

### CfDNA isolation, library preparation, and ctDNA capture

CfDNA was isolated from 1 to 1.5 mL plasma from each sample using the MagMax™ Cell-free DNA Isolation Kit (Thermo Fisher Scientific) according to the manufacturer’s instructions. Subsequently, the cfDNA was modified and amplified to prepare the multiplexed paired-end sequencing libraries with the dual index using GenTrack Library Preparation Kit (GenScript). The sequencing libraries were prepared by end-repair, A-tailing, adapter ligation, and library amplification, with 7 to 14 cycles of PCR using the Phusion High-Fidelity PCR Master Mix (Thermo Fisher Scientific). The ctDNA yield was quantified with the Qubit™ dsDNA HS Assay Kit (Thermo Fisher Scientific). For hybrid capturing of selected ctDNAs, a hybrid probe library with 6394 probes was designed and synthesized by GenScript. Magnetic bead-based capturing, washing, and elution were conducted according to the manufacturer’s protocol. The post-capture PCR amplification was set as eight cycles. Preliminary quantification of the captured ctDNA libraries was conducted using the Qubit™ dsDNA HS Assay Kit. The qualified libraries, ranging from 130 bp to 170 bp in length, were analyzed by the Agilent High Sensitivity DNA Analysis Kit, and subjected to 150 base paired-end sequencing on the Illumina NovaSeq 6000 instrument.

### Bioinformatics pipeline for the process of ctDNA profiling data

The ctDNA/cfDNA data from the hybrid capturing sequencing of patients with NPC, and the public datasets of the patients with prostate cancer (PRJNA554329, SRA database) and the healthy cohorts (SRP147273, SRA database), were processed according to the following steps. QC of sequencing reads was performed by fastp software ^29^. Reads with Phred quality scores < 30 in > 20% bases were discarded and the clean reads were used for the following analysis. The reference genome of hg38, covering the autosomes and sex chromosomes, was downloaded from the UCSC Genome Browser, and the paired-end reads were mapped to the reference genome with the BWA mem function ^30^. To further evaluate the sequencing data quality, the on-target rate, the average depth of every exon in the Enrichers, and the DNA size features were analyzed by the BEDTools coverage ^31^, the SAMtools depth ^32^, and the manual shell script, respectively (Supplementary Table 5; Supplementary Figure 4). A manual shell script was used to perform size selection *in silico*, and DNA fragments with lengths of 130-170 bp were selected for ctDNA signal enrichment. The mutation calling pipeline was accessed by the GATK functions of MarkDuplicates, FixMateInformation, BaseRecalibrator, ApplyBQSR, and Mutect2 for mutation calling ^33^. Then, the raw mutations were annotated to refGene, cytoband, and clinvar libraries by ANNOVAR ^34^. The sequencing QC parameters of the allelic depth of the reference and alteration alleles (AD) and approximate read depth (DP) in Variant Call Format (VCF) files were considered. Mutations with the AD of altered alleles = 0 or DP = 0 were discarded. Next, the R package maftools was used to transform the VCF file to MAF format ^35^ and remove the silent mutations. To further reduce the interference of SNP, we analyzed the SNP potential of each filtered mutation using the population WBC data generated by the 1000 Genome Project (1000G) of 2504 healthy donors from different districts worldwide ^36^. The SNP reference file was established by collecting the variants in the 1000G datasets of the mutations with frequencies > 1% and the total number of alternate alleles in called genotypes (AC) > 1000. The mutation working list was set by removing the mutations involved in the SNP reference files, which further resorts to the subsequent mapping to the Enricher exons.

### Statistical analysis

Data analysis was performed using R software (Version 4.0.5) ^37^. The overall survival (OS) and progress-free survival (PFS) analysis was based on the R packages survival and survminer. The OS data were fit to a univariable Cox regression model and tested by a log-rank test. The Pearson’s correlation coefficient was used to measure the similarity of variables to tTMB. The receiver operating characteristic (ROC) curve was plotted with the assistance of the R package pROC ^38^. The threshold, AUC, specificity, and sensitivity are listed in Supplementary Table 6. The main visualization work was based on the R package ggplot2 ^39^. The R package GenomicRanges was used to deal with genomic problems ^40^. The two-sided Wilcoxon test and t-test were used to assess the statistical differences between groups unless mentioned otherwise. Different levels of statistical significance were assessed based on specific p-values (*P < 0.05, **P < 0.01, and ***P < 0.001).

## Data and code availability

All data for this study are included in this article and its supplemental materials. The hybrid-capturing sequencing data of the ctDNA in patients with NPC are available via NCBI SRA: PRJNA796715 (https://www.ncbi.nlm.nih.gov/bioproject/796715). The source code was mainly based on the R language. Custom scripts and additional data related to support the findings of this work will be available to the academic community upon reasonable request to the lead contact.

## Acknowledgments

We thank LetPub (www.letpub.com) for its linguistic assistance during the preparation of this manuscript.

## Author contributions

Lv.Z. conceived and designed the project and provided overall guidance. M.L. and Lv.Z. developed the algorithm, wrote the programs, and performed the computational analysis. M.L., S.X. and T.H. designed and equally contributed to conducting all the biological experiments. T.H. collected the plasma samples of patients with NPC. T.S., J.K. and C.L. assisted parts of experiments. Ling.Z. and X.L. provide valuable advises. All authors reviewed the manuscript and provided comments.

## Figure and table legends

### Figure legends

**Supplementary Figure 1. The patient capture rate of Enricher-v1 in the top 10 most lethal cancers worldwide. The cancer types with 90% of the patients who can be captured at least once with Enricher-v1 are highlighted in the green area**.

**Supplementary Figure 2. Evaluation of OS and PFS of patients with multiple cancers by MaGICv1 at Q3 threshold**.

**a**. The patients are classified as LMG (TMB = 0), MMG (1 ≤ KME ≤ Q3 value threshold), and HMG (TMB > Q3 value threshold) of each cancer. The P-value between each comparison is labeled in the heatmap and assigned to different colors (log-rank test).

**b**. The univariable Cox curves of PFS and OS of representative cancer type (kidney cancer and head and neck cancer) where the patients are classified by MaGICv1 (HMG thresholds: Q3 value).

OS, overall survival. PFS, progress-free survival. TMB, tumor mutation burden. LMG, low-mutation group. MMG, middle-mutation group. HMG, high-mutation group. Q3, upper quantile. NS, not significant. NA, data not available.

**Supplementary Figure 3. Performance of MaGICv1 in prognosis evaluation of the top 10 most lethal cancers worldwide in the context of tissue biopsy**.

The patients are classified as LMG (TMB = 0), MMG (1 ≤ KME ≤ median or Q3 value threshold), and HMG (TMB > median or Q3 value threshold) of each cancer. The P-value between each comparison is labeled in the heatmap and assigned to different colors (log-rank test).

OS, overall survival. PFS, progress-free survival. TMB, tumor mutation burden. LMG, low-mutation group. MMG, middle-mutation group. HMG, high-mutation group. Q3, upper quantile. NS, not significant. NA, data not available.

**Supplementary Figure 4. The quality control of ctDNA capture sequencing data from patients with nasopharyngeal carcinoma**

**a**. The on-target rate of the exons in Enricher-v1. The error bars in the barplot represent mean on-target rate ± standard deviation.

**b**. Histogram shows the fragment length of aligned ctDNA for sequencing. The bin width was set to 5bp.

**c**. Boxplot shows the fragment length of aligned ctDNA after deduplicated and size selection within each sample.

ctDNA, circulating tumor DNA.

**Supplementary Figure 5. The NMR and CT images of patients with nasopharyngeal carcinoma at the time points before-FRC and after-SRC**.

NMR, nuclear magnetic resonance map. CT, computerized tomography. FRC, first-round chemotherapy. SRC, second round chemotherapy.

**Supplementary Figure 6. Evaluation of OS and PFS of patients with multiple cancers by MaGICv2 at Q3 threshold**.

**a**. The patients are classified as LMG (KME = 0), MMG (1 ≤ KME ≤ Q3 value threshold), and HMG (KME > Q3 value threshold) of each cancer. The P-value between each comparison is labeled in the heatmap and assigned to different colors (log-rank test).

**b**. The univariable Cox curves of PFS and OS of representative cancer types (head and neck cancer and brain cancer) where the patients are classified by MaGICv2 (HMG thresholds: Q3 value).

OS, overall survival. PFS, progress-free survival. TMB, tumor mutation burden. mVAF, mean variant allele frequency. KME, the number of key mutated exons. LMG, low-mutation group. MMG, middle-mutation group. HMG, high-mutation group. Q3, the upper quantile. NS, not significant. NA, data not available.

**Supplementary Figure 7. Performance of MaGICv2 in prognosis evaluation of the top 10 most lethal cancers worldwide in the context of tissue biopsy**

OS, overall survival. PFS, progress-free survival. TMB, tumor mutation burden. mVAF, mean variant allele frequency. KME, the number of key mutated exons. LMG, low-mutation group. MMG, middle-mutation group. HMG, high-mutation group. Q3, the upper quantile. NS, not significant. NA, data not available.

**Supplementary Figure 8. Comparison of the performance of the two versions of MaGICs in prognosis evaluation (HMG thresholds: Q3 value)**.

OS, overall survival. PFS, progress-free survival. TMB, tumor mutation burden. LMG, low-mutation group. MMG, middle-mutation group. HMG, high-mutation group. Q3, upper quantile. NS, not significant. NA, data not available.

**Supplementary Figure 9. Comparison of the performance of the two versions of MaGICs in prognosis evaluation of the top 10 most lethal cancers worldwide**.

OS, overall survival. PFS, progress-free survival. TMB, tumor mutation burden. LMG, low-mutation group. MMG, middle-mutation group. HMG, high-mutation group. Q3, upper quantile. NS, not significant. NA, data not available.

**Supplementary Figure 10. Comparison of the performance of MaGIC-v2 with other published panels in combination with KME measurement in prognosis evaluation of the top 10 most lethal cancers worldwide**.

LMG, low-mutation group. MMG, middle-mutation group. HMG, high-mutation group. KME, the number of key mutated exons.

**Supplementary Figure 11. Comparison of the performance of MaGIC-v2 with MaGIC-v1 and other different combinations of Enrichers and measurements in chemosensitivity prediction of nasopharyngeal carcinoma**.

bTMB, blood-based tumor mutation burden. mVAF, mean variant allele frequency. FRC, first-round chemotherapy. SRC, second round chemotherapy. a-b, the difference value of after-FRC and before-FRC. ROC, receiver operation characteristic. AUC, the area under curve.

### Table legends

**Supplementary Table 1. TCGA whole exome sequencing projects used for tissue biopsy analysis**

**Supplementary Table 2. The Enricher-v1 targeted exons**.

**Supplementary Table 3. The Enricher-v2 targeted exons**.

**Supplementary Table 4. The characteristics of patients with nasopharyngeal carcinoma**.

**Supplementary Table 5. Statistical analysis of the quality of sequencing data**.

**Supplementary Table 6 Data of the ROC curves in ctDNA analysis**.

## References

1 Mohan, S. et al. Profiling of Circulating Free DNA Using Targeted and Genome-wide Sequencing in Patients with SCLC. J Thorac Oncol 15, 216–230, doi:10.1016/j.jtho.2019.10.007 (2020).

2 Im, Y. R., Tsui, D. W. Y., Diaz, L. A., Jr. & Wan, J. C. M. Next-Generation Liquid Biopsies: Embracing Data Science in Oncology. Trends Cancer 7, 283–292, doi:10.1016/j.trecan.2020.11.001 (2021).

3 Dawson, S. J. et al. Analysis of circulating tumor DNA to monitor metastatic breast cancer. N Engl J Med 368, 1199–1209, doi:10.1056/NEJMoa1213261 (2013).

4 Gandara, D. R. et al. Blood-based tumor mutational burden as a predictor of clinical benefit in non-small-cell lung cancer patients treated with atezolizumab. Nat Med 24, 1441–1448, doi:10.1038/s41591-018-0134-3 (2018).

5 Fujii, Y. et al. Identification and monitoring of mutations in circulating cell-free tumor DNA in hepatocellular carcinoma treated with lenvatinib. J Exp Clin Cancer Res 40, 215, doi:10.1186/s13046-021-02016-3 (2021).

6 Crowley, E., Di Nicolantonio, F., Loupakis, F. & Bardelli, A. Liquid biopsy: monitoring cancer-genetics in the blood. Nat Rev Clin Oncol 10, 472–484, doi:10.1038/nrclinonc.2013.110 (2013).

7 Garcia-Murillas, I. et al. Mutation tracking in circulating tumor DNA predicts relapse in early breast cancer. Sci Transl Med 7, 302ra133, doi:10.1126/scitranslmed.aab0021 (2015).

8 Tie, J. et al. Circulating tumor DNA analysis detects minimal residual disease and predicts recurrence in patients with stage II colon cancer. Sci Transl Med 8, 346ra392, doi:10.1126/scitranslmed.aaf6219 (2016).

9 Lui, Y. Y. et al. Predominant hematopoietic origin of cell-free DNA in plasma and serum after sex-mismatched bone marrow transplantation. Clin Chem 48, 421–427 (2002).

10 Xia, L. et al. Statistical analysis of mutant allele frequency level of circulating cell-free DNA and blood cells in healthy individuals. Sci Rep 7, 7526, doi:10.1038/s41598-017-06106-1 (2017).

11 Yong, E. Cancer biomarkers: Written in blood. Nature 511, 524–526, doi:10.1038/511524a (2014).

12 Murtaza, M. et al. Non-invasive analysis of acquired resistance to cancer therapy by sequencing of plasma DNA. Nature 497, 108–112, doi:10.1038/nature12065 (2013).

13 Chaudhuri, A. A. et al. Early Detection of Molecular Residual Disease in Localized Lung Cancer by Circulating Tumor DNA Profiling. Cancer Discov 7, 1394–1403, doi:10.1158/2159-8290.CD-17-0716 (2017).

14 Abbosh, C. et al. Phylogenetic ctDNA analysis depicts early-stage lung cancer evolution. Nature 545, 446–451, doi:10.1038/nature22364 (2017).

15 Phallen, J. et al. Early Noninvasive Detection of Response to Targeted Therapy in Non-Small Cell Lung Cancer. Cancer Res 79, 1204–1213, doi:10.1158/0008-5472.CAN-18-1082 (2019).

16 Hu, Y. et al. False-Positive Plasma Genotyping Due to Clonal Hematopoiesis. Clin Cancer Res 24, 4437–4443, doi:10.1158/1078-0432.CCR-18-0143 (2018).

17 Leal, A. et al. White blood cell and cell-free DNA analyses for detection of residual disease in gastric cancer. Nat Commun 11, 525, doi:10.1038/s41467-020-14310-3 (2020).

18 Newman, A. M. et al. An ultrasensitive method for quantitating circulating tumor DNA with broad patient coverage. Nat Med 20, 548–554, doi:10.1038/nm.3519 (2014).

19 Newman, A. M. et al. Integrated digital error suppression for improved detection of circulating tumor DNA. Nat Biotechnol 34, 547–555, doi:10.1038/nbt.3520 (2016).

20 Razavi, P. et al. High-intensity sequencing reveals the sources of plasma circulating cell-free DNA variants. Nat Med 25, 1928–1937, doi:10.1038/s41591-019-0652-7 (2019).

21 Teo, Y. V. et al. Cell-free DNA as a biomarker of aging. Aging Cell 18, e12890, doi:10.1111/acel.12890 (2019).

22 Ramesh, N. et al. Decoding the evolutionary response to prostate cancer therapy by plasma genome sequencing. Genome Biol 21, 162, doi:10.1186/s13059-020-02045-9 (2020).

23 Burgener, J. M. et al. Tumor-Naive Multimodal Profiling of Circulating Tumor DNA in Head and Neck Squamous Cell Carcinoma. Clin Cancer Res 27, 4230–4244, doi:10.1158/1078-0432.CCR-21-0110 (2021).

24 Cheng, D. T. et al. Memorial Sloan Kettering-Integrated Mutation Profiling of Actionable Cancer Targets (MSK-IMPACT): A Hybridization Capture-Based Next-Generation Sequencing Clinical Assay for Solid Tumor Molecular Oncology. J Mol Diagn 17, 251–264, doi:10.1016/j.jmoldx.2014.12.006 (2015).

25 Murphy, D. Comprehensive Analyses of Circulating Cell-Free Tumor DNA. 28.

26 Wang, Z. et al. Assessment of Blood Tumor Mutational Burden as a Potential Biomarker for Immunotherapy in Patients With Non-Small Cell Lung Cancer With Use of a Next-Generation Sequencing Cancer Gene Panel. JAMA Oncol 5, 696–702, doi:10.1001/jamaoncol.2018.7098 (2019).

27 Kim, S. T. et al. Prospective Feasibility Study for Using Cell-Free Circulating Tumor DNA-Guided Therapy in Refractory Metastatic Solid Cancers: An Interim Analysis. JCO Precis Oncol 1, 1–15, doi:10.1200/PO.16.00059 (2017).

28 Sharaf, R. et al. FoundationOne CDx testing accurately determines whole arm 1p19q codeletion status in gliomas. Neurooncol Adv 3, vdab017, doi:10.1093/noajnl/vdab017 (2021).

29 Chen, S., Zhou, Y., Chen, Y. & Gu, J. fastp: an ultra-fast all-in-one FASTQ preprocessor. Bioinformatics 34, i884–i890, doi:10.1093/bioinformatics/bty560 (2018).

30 Li, H. & Durbin, R. Fast and accurate short read alignment with Burrows-Wheeler transform. Bioinformatics 25, 1754–1760, doi:10.1093/bioinformatics/btp324 (2009).

31 Quinlan, A. R. & Hall, I. M. BEDTools: a flexible suite of utilities for comparing genomic features. Bioinformatics 26, 841–842, doi:10.1093/bioinformatics/btq033 (2010).

32 Li, H. et al. The Sequence Alignment/Map format and SAMtools. Bioinformatics 25, 2078–2079, doi:10.1093/bioinformatics/btp352 (2009).

33 McKenna, A. et al. The Genome Analysis Toolkit: a MapReduce framework for analyzing next-generation DNA sequencing data. Genome Res 20, 1297–1303, doi:10.1101/gr.107524.110 (2010).

34 Venables, W. N. & Ripley, B. D. Modern Applied Statistics with S. Fourth edn, (Springer, 2002).

35 Mayakonda, A., Lin, D. C., Assenov, Y., Plass, C. & Koeffler, H. P. Maftools: efficient and comprehensive analysis of somatic variants in cancer. Genome Res 28, 1747–1756, doi:10.1101/gr.239244.118 (2018).

36 Genomes Project, C. et al. A global reference for human genetic variation. Nature 526, 68–74, doi:10.1038/nature15393 (2015).

37 Team, R. C. R: A Language and Environment for Statistical Computing. (R Foundation for Statistical Computing, 2020).

38 Robin, X. et al. pROC: an open-source package for R and S+ to analyze and compare ROC curves. BMC Bioinformatics 12, 77, doi:10.1186/1471-2105-12-77 (2011).

39 Wickham, H. ggplot2: Elegant Graphics for Data Analysis. (Springer-Verlag New York, 2016).

40 Lawrence, M. et al. Software for computing and annotating genomic ranges. PLoS Comput Biol 9, e1003118, doi:10.1371/journal.pcbi.1003118 (2013).

41 Sung, H. et al. Global Cancer Statistics 2020: GLOBOCAN Estimates of Incidence and Mortality Worldwide for 36 Cancers in 185 Countries. CA Cancer J Clin 71, 209–249, doi:10.3322/caac.21660 (2021).

42 Geurts, M. & van den Bent, M. J. Hypermutated recurrences: Identifying the clinical relevance. Neuro Oncol 23, 1805–1806, doi:10.1093/neuonc/noab192 (2021).

43 Martínez-Jiménez, F. et al. A compendium of mutational cancer driver genes. Nature reviews. Cancer 20, 555–572, doi:10.1038/s41568-020-0290-x (2020).

44 Phallen, J. et al. Direct detection of early-stage cancers using circulating tumor DNA. Sci Transl Med 9, doi:10.1126/scitranslmed.aan2415 (2017).

45 Jaiswal, S. & Ebert, B. L. Clonal hematopoiesis in human aging and disease. Science 366, eaan4673, doi:10.1126/science.aan4673 (2019).

46 Abbosh, C., Swanton, C. & Birkbak, N. J. Clonal haematopoiesis: a source of biological noise in cell-free DNA analyses. Ann Oncol 30, 358–359, doi:10.1093/annonc/mdy552 (2019).

