## Supplementary materials for "Circulating tumor DNA profiling approach based on in silico background elimination guides patient classification of multiple cancers"

**This PDF file includes:**

Figures S1 to S11
Data S1 to S6

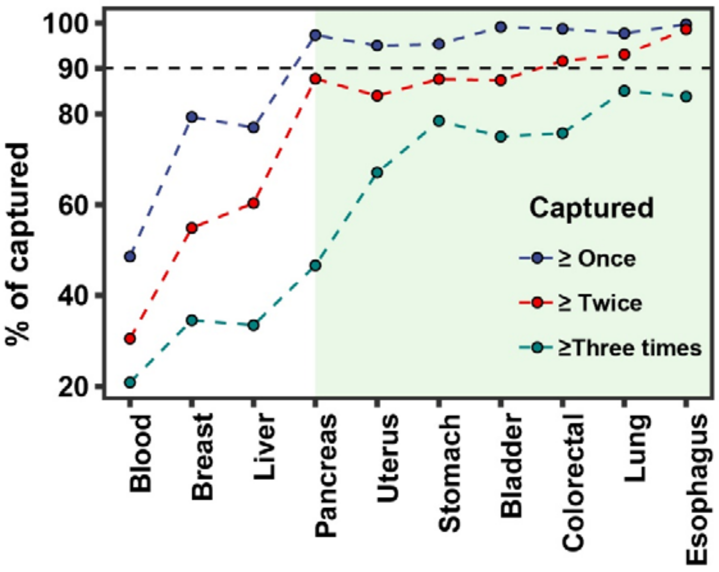


**Figureure. S1.**

**The patient capture rate of Enricher-v1 in the top 10 most lethal cancers worldwide.** The cancer types with 90% of the patients who can be captured at least once with Enricher-v1 are highlighted in the green area.

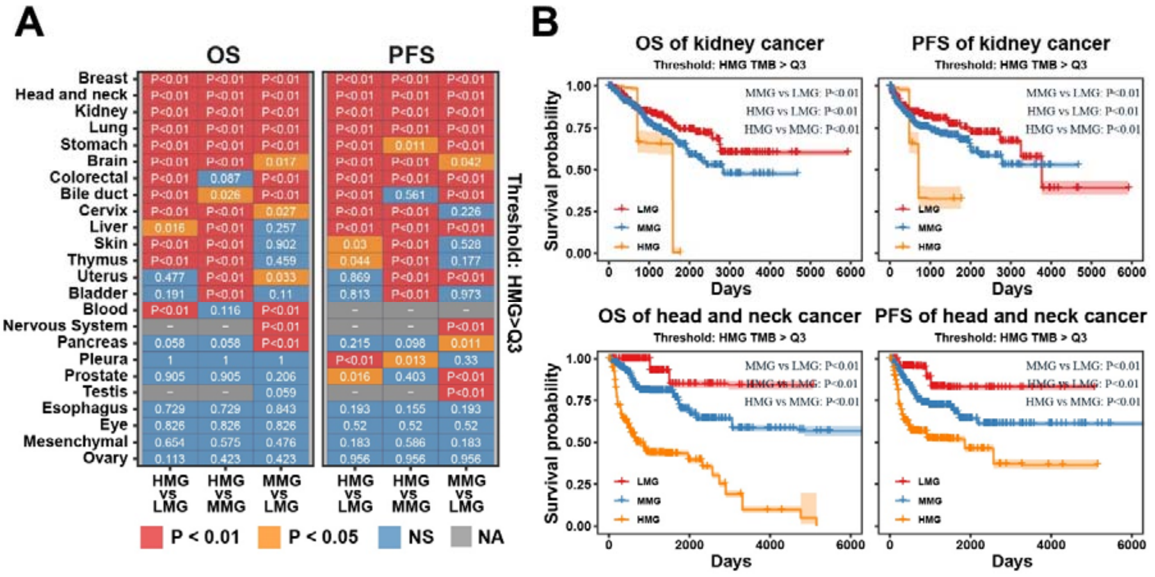


**Figure. S2.**

**Evaluation of OS and PFS of patients with multiple cancers by MaGICv1 at Q3 threshold.** (**A**), The patients are classified as LMG (TMB = 0), MMG (1 ≤ KME ≤ Q3 value threshold), and HMG (TMB > Q3 value threshold) of each cancer. The P-value between each comparison is labeled in the heatmap and assigned to different colors (log-rank test). (**B**), The univariable Cox curves of PFS and OS of representative cancer type (kidney cancer and head and neck cancer) where the patients are classified by MaGICv1 (HMG thresholds: Q3 value). OS, overall survival. PFS, progress-free survival. TMB, tumor mutation burden. LMG, low-mutation group. MMG, middle-mutation group. HMG, high-mutation group. Q3, upper quantile. NS, not significant. NA, data not available.

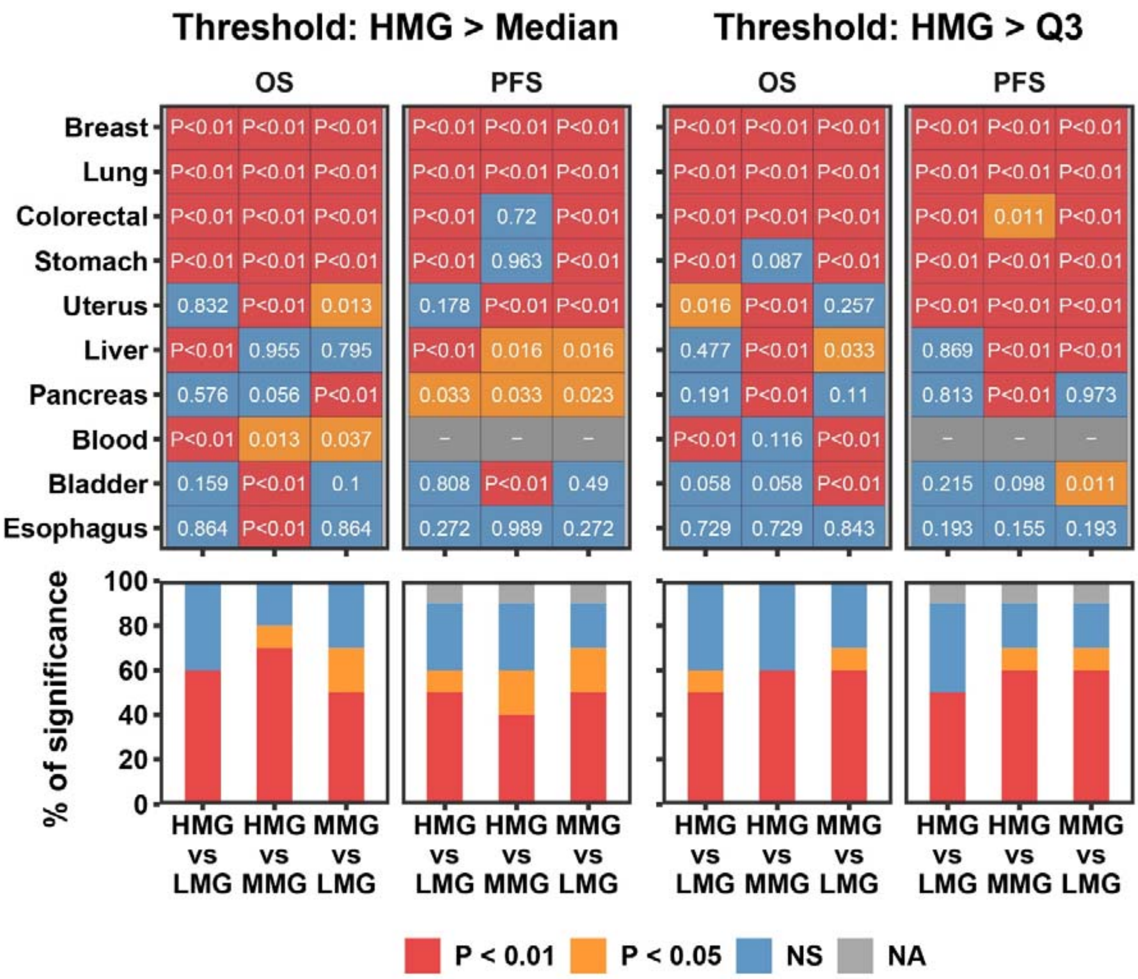


**Figure. S3.**

**Performance of MaGICv1 in prognosis evaluation of the top 10 most lethal cancers worldwide in the context of tissue biopsy.** The patients are classified as LMG (TMB = 0), MMG (1 ⩽ KME ⩽ median or Q3 value threshold), and HMG (TMB > median or Q3 value threshold) of each cancer. The P-value between each comparison is labeled in the heatmap and assigned to different colors (log-rank test). OS, overall survival. PFS, progress-free survival. TMB, tumor mutation burden. LMG, low-mutation group. MMG, middle-mutation group. HMG, high-mutation group. Q3, upper quantile. NS, not significant. NA, data not available.

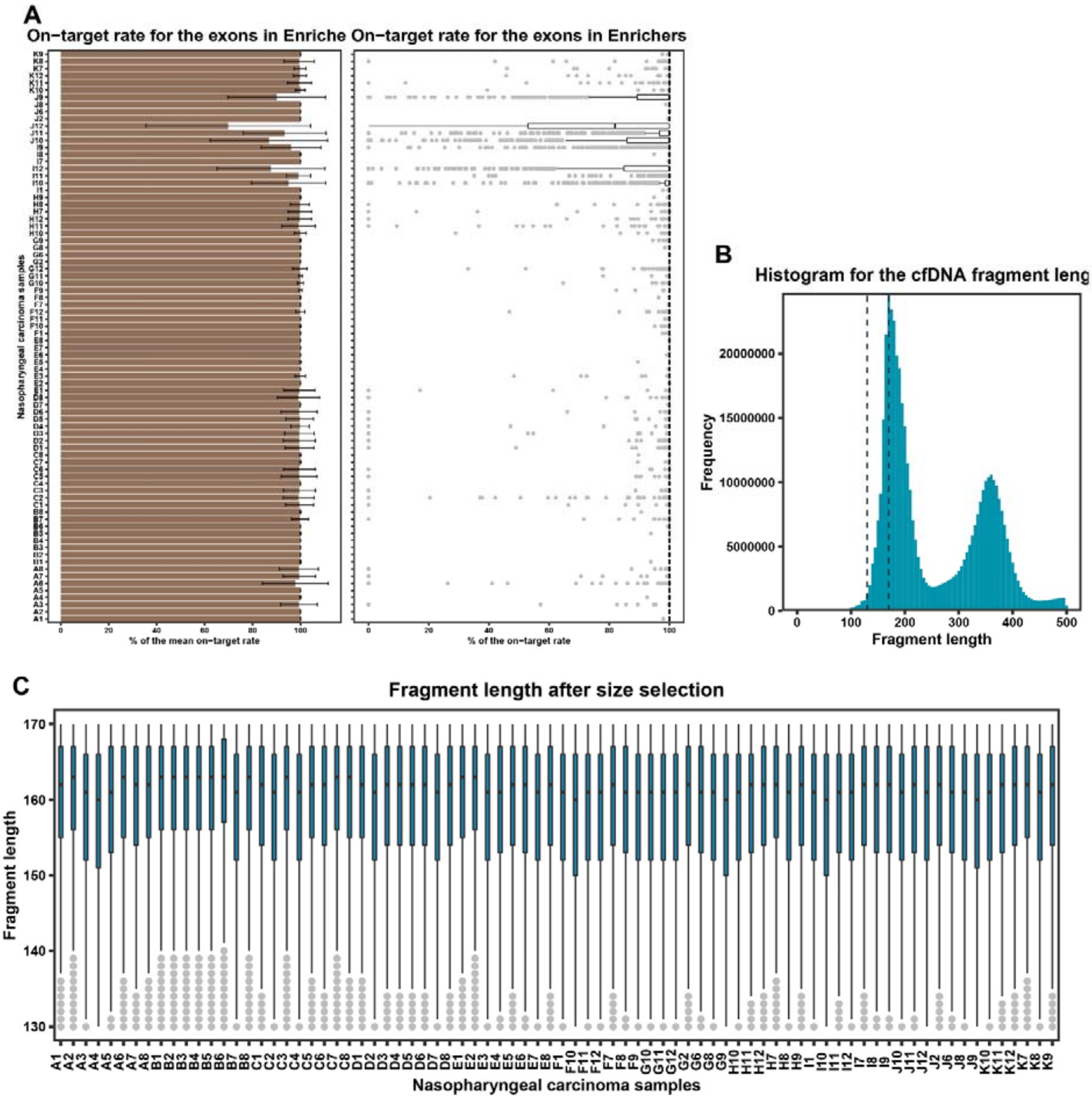


**Figure. S4.**

**The quality control of ctDNA capture sequencing data from patients with nasopharyngeal carcinoma.** (**A**), The on-target rate of the exons in Enricher-v1. The error bars in the barplot represent mean on-target rate ± standard deviation. (**B**), Histogram shows the fragment length of aligned ctDNA for sequencing. The bin width was set to 5bp. (**C**), Boxplot shows the fragment length of aligned ctDNA after deduplicated and size selection within each sample. ctDNA, circulating tumor DNA.

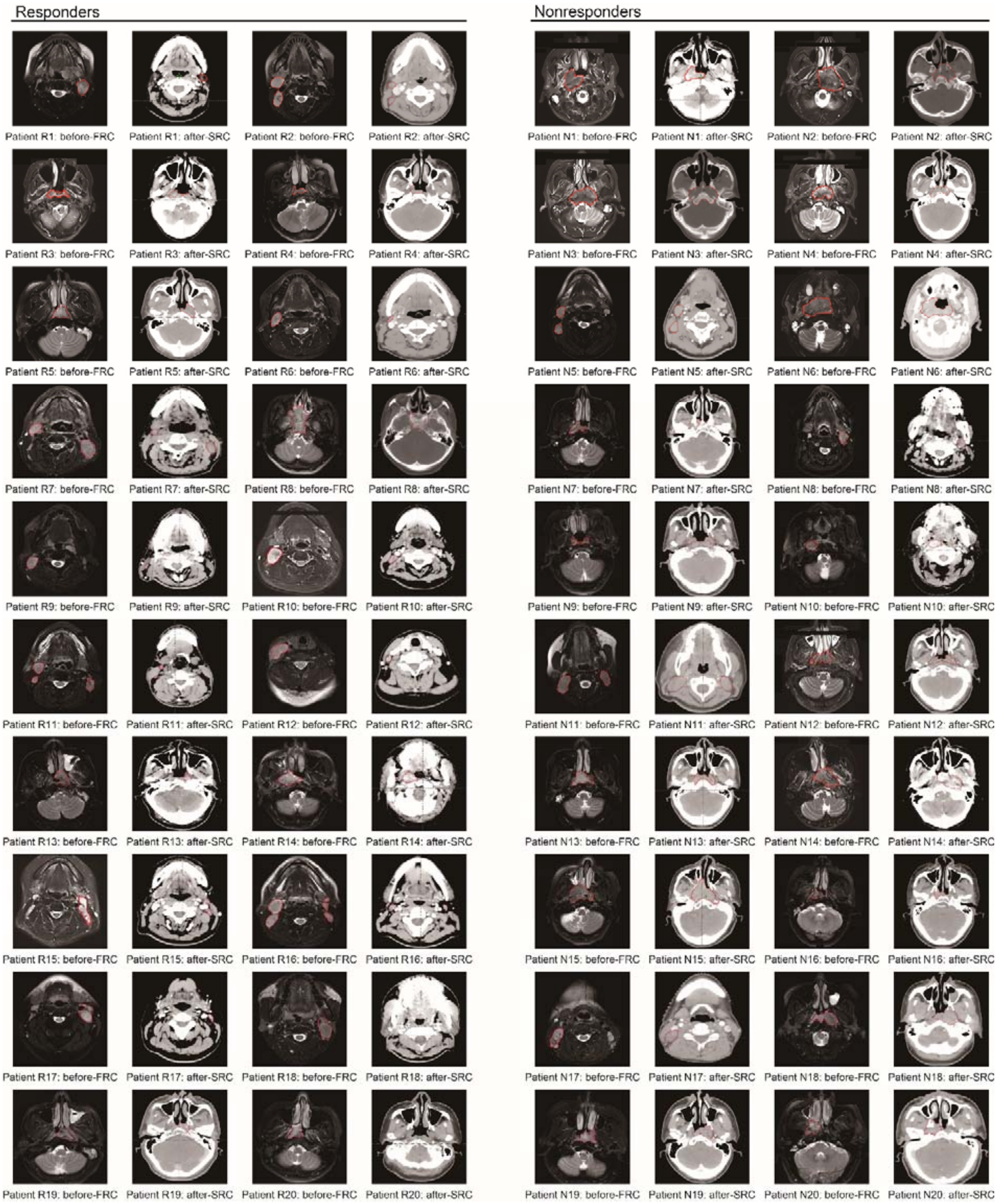


**Figure. S5.**

**The NMR and CT images of patients with nasopharyngeal carcinoma at the time points before-FRC and after-SRC.** NMR, nuclear magnetic resonance map. CT, computerized tomography. FRC, first-round chemotherapy. SRC, second round chemotherapy.

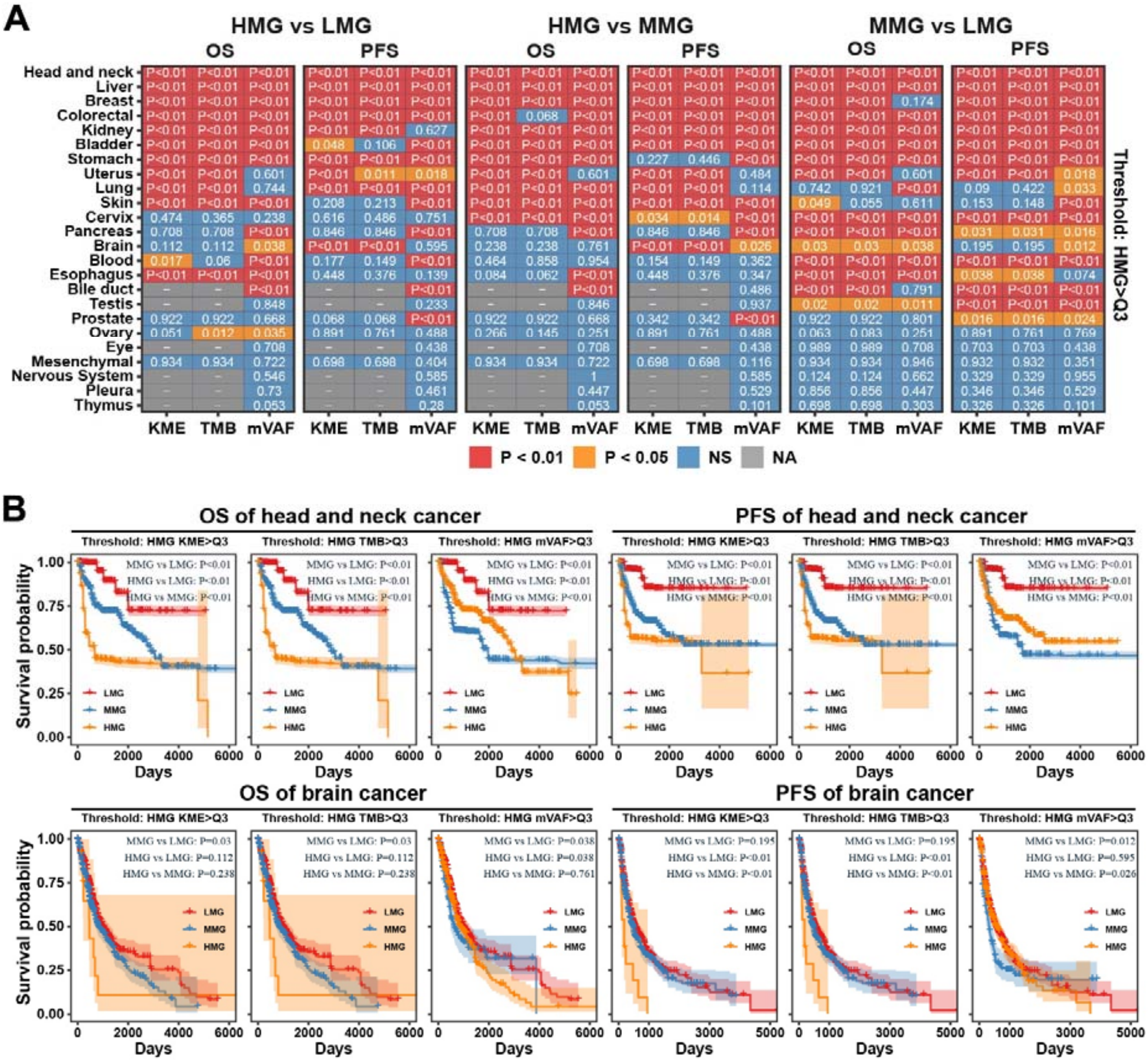


**Figure. S6.**

**Evaluation of OS and PFS of patients with multiple cancers by MaGICv2 at Q3 threshold.** (**A**), The patients are classified as LMG (KME = 0), MMG (1 ≤ KME ≤ Q3 value threshold), and HMG (KME > Q3 value threshold) of each cancer. The P-value between each comparison is labeled in the heatmap and assigned to different colors (log-rank test). (**B**), The univariable Cox curves of PFS and OS of representative cancer types (head and neck cancer and brain cancer) where the patients are classified by MaGICv2 (HMG thresholds: Q3 value). OS, overall survival. PFS, progress-free survival. TMB, tumor mutation burden. mVAF, mean variant allele frequency. KME, the number of key mutated exons. LMG, low-mutation group. MMG, middle-mutation group. HMG, high-mutation group. Q3, the upper quantile. NS, not significant. NA, data not available.

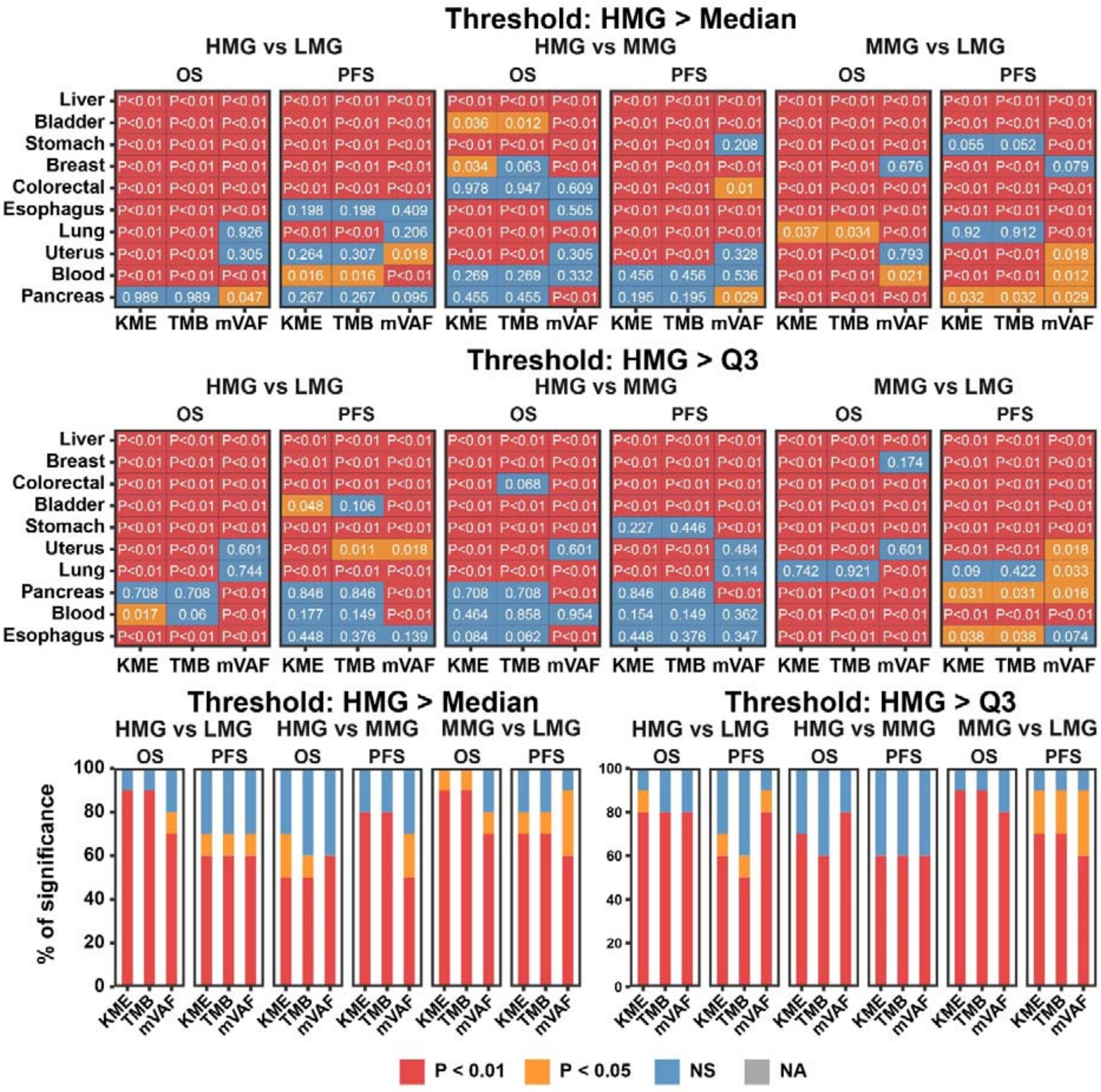


**Figure. S7.**
**Performance of MaGICv2 in prognosis evaluation of the top 10 most lethal cancers worldwide in the context of tissue biopsy.** OS, overall survival. PFS, progress-free survival. TMB, tumor mutation burden. mVAF, mean variant allele frequency. KME, the number of key mutated exons. LMG, low-mutation group. MMG, middle-mutation group. HMG, high-mutation group. Q3, the upper quantile. NS, not significant. NA, data not available.

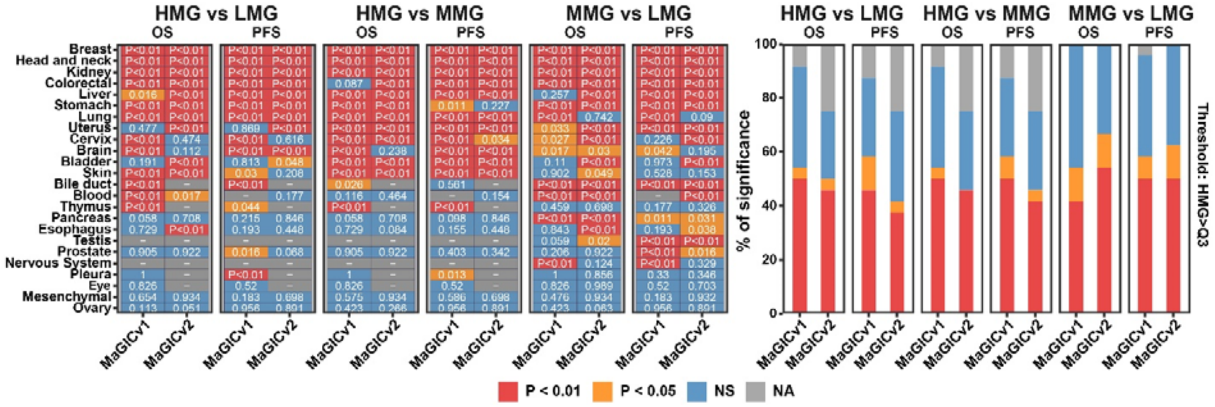


**Figure. S8.**

**Comparison of the performance of the two versions of MaGICs in prognosis evaluation (HMG thresholds: Q3 value).** OS, overall survival. PFS, progress-free survival. TMB, tumor mutation burden. LMG, low-mutation group. MMG, middle-mutation group. HMG, high-mutation group. Q3, upper quantile. NS, not significant. NA, data not available.

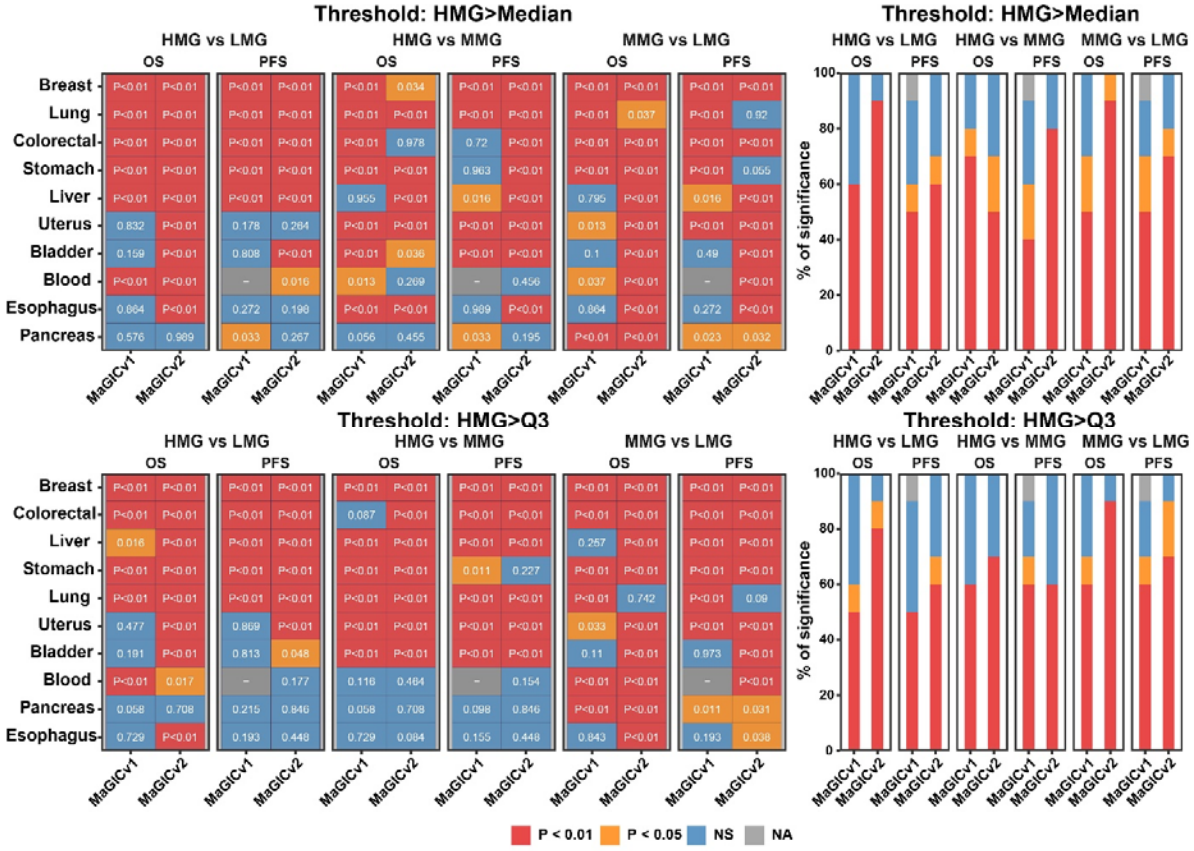


**Figure. S9.**
**Comparison of the performance of the two versions of MaGICs in prognosis evaluation of the top 10 most lethal cancers worldwide.** OS, overall survival. PFS, progress-free survival. TMB, tumor mutation burden. LMG, low-mutation group. MMG, middle-mutation group. HMG, high-mutation group. Q3, upper quantile. NS, not significant. NA, data not available.

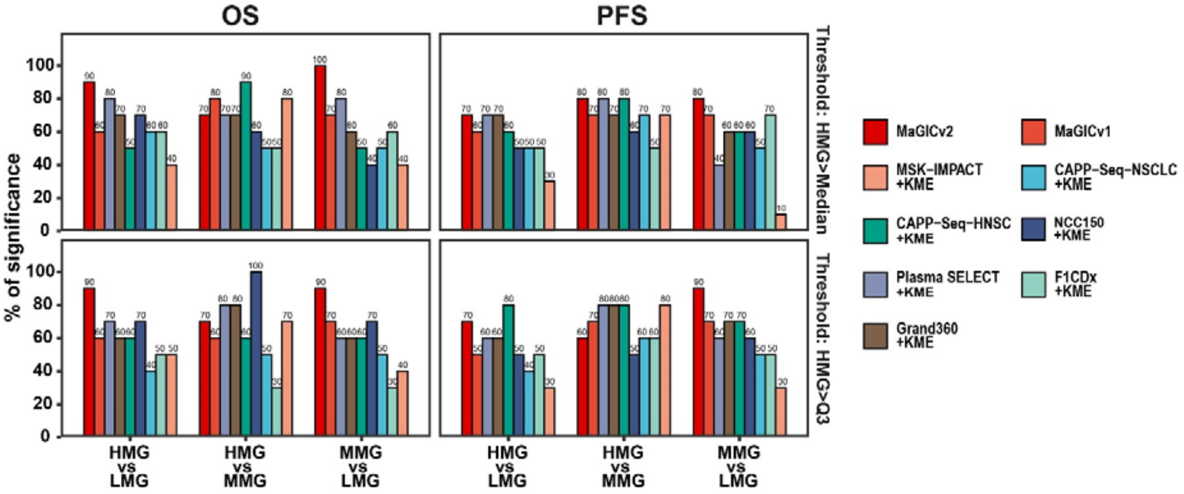


**Figure. S10.**
**Comparison of the performance of MaGIC-v2 with other published panels in combination with KME measurement in prognosis evaluation of the top 10 most lethal cancers worldwide.** LMG, low-mutation group. MMG, middle-mutation group. HMG, high-mutation group. KME, the number of key mutated exons.

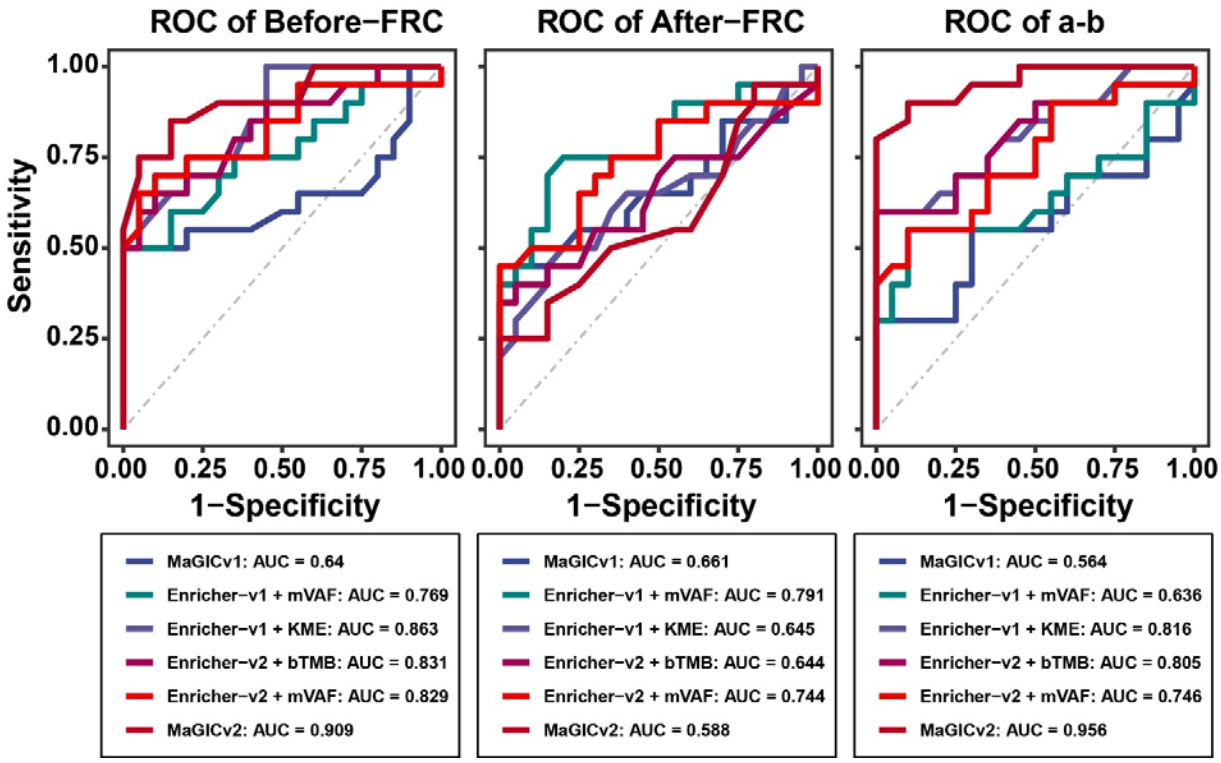


**Figure. S11.**
**Comparison of the performance of MaGIC-v2 with MaGIC-v1 and other different combinations of Enrichers and measurements in chemosensitivity prediction of nasopharyngeal carcinoma.** bTMB, blood-based tumor mutation burden. mVAF, mean variant allele frequency. FRC, first-round chemotherapy. SRC, second round chemotherapy. a-b, the difference value of after-FRC and before-FRC. ROC, receiver operation characteristic. AUC, the area under curve.

**Data S1.**
TCGA whole exome sequencing projects used for tissue biopsy analysis.

**Data S2.**
The Enricher-v1 targeted exons.

**Data S3.**
The Enricher-v2 targeted exons.

**Data S4.**
The characteristics of patients with nasopharyngeal carcinoma.

**Data S5.**
Statistical analysis of the quality of sequencing data.

**Data S6.**
Data of the ROC curves in ctDNA analysis.
